## Supplementary Material for "Single-Molecule RNA Sizing Enables Quantitative Analysis of Alternative Transcription Termination"

This PDF file includes:

→ Materials and Methods
→ Supplementary Figures 1 to 29
→ Supplementary Tables 1 to 8

### Materials and Methods

#### Materials

Commercial reagents implemented in this work included nuclease-free water (Ambion, catalog number AM9937), 100 × Tris-EDTA buffer solution concentrate (Sigma-Aldrich, Catalog number T9285), Lithium chloride for molecular biology ≥ 99% purity (Sigma-Aldrich, Catalog number L9650), Tris-HCl BioPerformance certified, ≥ 99% purity (Sigma-Aldrich, catalog number T5941). Buffer solutions prepared from these reagents were filtered with 0.22 µm Millipore syringe filter units (MF-Merck Millipore™, Catalog number GSWP04700).

For enzymatic reactions the following products were used: DraIII-HF (20,000 units/mL, New England Biolabs (NEB), Catalog number R3510S), ScaI-HF (20,000 units/mL, NEB, Catalog number R3122S), *Escherichia coli* Topoisomerase I (5,000 units/mL, NEB, Catalog Number M0301S), DNase I (2,000 units/mL, NEB, Catalog Number M0303S), HiScribe™ T7 Quick High Yield RNA Synthesis Kit (NEB, Catalog number E2050S).

DNA and RNA purification were performed with the following purification kits: Monarch PCR & DNA Clean-up Kit (5 µg, NEB, Catalog number T1030S) for DNA, and Monarch RNA Cleanup Kit (50 µg, New England Biolabs, Catalg Number T2040S) for RNA.

For this study we used Labcon Eclipse™ 10 µL Graduated Pipette Tips with UltraFine™ Point (Thermo Fisher Scientific, Catalog number 16603912), Eppendorf DNA LoBind® Tubes (Thermo Fisher Scientific, Catalog number 10686313 and 107008704), 0.2 mL RNase-free PCR tubes (Thermo Fisher Scientific, Catalog number AM12225), and Invitrogen RNaseZap™ RNase Decontamination Solution (Thermo Fisher Scientific, Catalog number AM9780).

For fabrication of glass nanopores, we used glass quartz capillaries with 0.2 mm inner diameter and 0.5 outer diameter (Sutter Instrument Company). For chip fabrication we used Sylgard 184 PDMS (Dow Corning, Catalog number 101697), microscope slides clear ground 1.0 – 1.2 mm (Thermo Fisher Scientific, Catalog number 1238-3118).

#### Circular DNA plasmid propagation

JM110 strain of *Escherichia coli* was used for propagation of circular plasmid with CTG repeats. 100 ng of plasmid was mixed with 50 µL of competent cells (0,085 M CaCl2, 15% glycerol). The mixture was incubated on ice for 30 minutes pror to heat shock at 42 °C for 90 sec. The cells were turned back on ice for 5 minutes and spread on LB agar plates supplemented with ampicillin (0,1 mg/ml). The plates were incubated at 37 °C overnight. Single colonies were picked and propagated in 50 ml of LB medium supplemented with ampicillin (0,1 mg/ml) at 37 °C overnight. Bacterial pellet was obtained by centrifugation at 4,000 rpm for 20 minutes. Plasmid DNA was extracted from the pellet using GeneJET Plasmid Miniprep Kit (Thermo Fisher, Catalog number K0503) according to the manufacturer's recommendations with modifications for larger volumes of starting bacterial culture. Plasmid DNA was eluted in 80 µL of nuclease-free water. DNA concentration was measured using Qubit™ dsDNA BR Assay Kit (Thermo Fisher Scientific, Catalog number Q32851) according to the manufacturer's recommendations.

**Circular DNA plasmid digestion: DraIII-HF**

DraIII-HF (20,000 units/mL, New England Biolabs (NEB), Catalog number R3510S) was used to linearize the circular DNA plasmids (plasmid sequence in Supplementary Table 1). Reaction of 2,000 ng of DNA was prepared according to the manufacturer's recommendations. In a 50 µL reaction, 2,000 ng of circular DNA plasmid were mixed with 2 µL of DraIII-HF (40 units), 5 µL of 10X rCutSmart Buffer, and nuclease-free water to achieve final reaction volume. The reaction components were mixed by pipetting and spin down for a couple of seconds. Reaction was incubated at 37 °C for 1 hour. Enzymes were kept on ice throughout the whole preparation procedure.

**Circular DNA plasmid digestion: ScaI-HF**

ScaI-HF (20,000 units/mL, NEB, Catalog number R3122S) was used to linearize circular DNA plasmid for Supplementary Figure 24 (plasmid sequence in Supplementary Table 5). Reaction of 2,000 ng of DNA was prepared according to the manufacturer's recommendations. In a 50 µL reaction, 2,000 ng of circular DNA plasmid were mixed with 2 µL of ScaI-HF (40 units), 5 µL of 10X rCutSmart Buffer, and nuclease-free water to achieve final reaction volume. The reaction components were mixed by pipetting and spin down for a couple of seconds. Reaction was incubated at 37 °C for 1 hour. Enzymes were kept on ice throughout the whole preparation procedure.

**Supercoiled plasmid relaxation: Topoisomerase I**

*Escherichia coli* Topoisomerase I (5,000 units/mL, NEB, Catalog Number M0301S) was used to relax a supercoiled circular DNA plasmid (plasmid sequence in Supplementary Table 1).

Reaction of 2,000 ng of DNA was prepared according to the manufacturer's recommendation. In a 100 µL reaction, 2,000 ng of circular DNA plasmid were mixed with 3 µL of Topoisomerase I (15 units), 10 µL of 10X rCutSmart Buffer, and nuclease-free water to achieve final reaction volume. The reaction components were mixed by pipetting and spin down for a couple of seconds. Reaction was incubated at 37 °C for 1 hour followed by incubation at 65 °C for 20 minutes to inactivate the enzyme. Enzymes were kept on ice throughout the whole preparation procedure.

**DNA purification**

Linearized or relaxed DNA was purified using Monarch PCR & DNA Clean-up Kit (5 µg, NEB, Catalog number T1030S). Binding and washing of the sample to the purification columns were performed as suggested by the manufacturer, following the suggested centrifugation protocols. After washing, the DNA was eluted with preheated (50 °C) nuclease-free water. The elution step was performed twice with 10 µL of nuclease-free water after 5 minutes incubation at room temperature to increase DNA yield. The concentration of DNA was estimated using a NanoDrop spectrophotometer.

***In vitro* transcription: T7 RNA polymerase**

Purified linear DNA and relaxed circular DNA were both *in vitro* transcribed using HiScribe™ T7 Quick High Yield RNA Synthesis Kit (NEB, Catalog number E2050S). For each, a 240 ng reaction was prepared according to the manufacturer's recommendation. In a 20 µL reaction, 240 ng of purified DNA were mixed with 2 µL of T7 RNA Polymerase (T7RNAP) Mix, 10 µL of NTP buffer mix (to achieve10 mM concentration of each NTP), and nuclease-free water to achieve final reaction volume. The reaction components were mixed by pipetting and spin down for a couple of seconds. The reaction was incubated at 37 °C for 4 hours. Variability in the yield of RNA synthesis (up to two orders of magnitude) was observed between different lots of the same T7 RNA polymerase Mix. We attribute this to variability in the concentration of the active enzyme between lots. Yield could be adjusted by varying the concentration of T7 RNA polymerase Mix in the reaction mixture, and yield was consistent within lots. Enzymes were kept on ice throughout the whole preparation procedure.

**DNA removal: DNase I treatment**

Transcription products were treated with DNase I (2,000 units/mL, NEB, Catalog Number M0303S) to remove DNA templates. 68 µL reaction mixture was prepared by adding 2 µL (4 units) of DNase I and 46 µL of nuclease-free water to the full volume (20 µL) of transcription products. The reaction components were mixed by pipetting and spin down for a couple of seconds. The reaction mixture was incubated at 37 °C for 15 minutes. Enzymes were kept on ice throughout the whole preparation procedure.

**RNA purification**

After DNA removal, RNA was purified using Monarch RNA Cleanup Kit (50 µg, New England Biolabs, Catalg Number T2040S) following the manufacturer's instructions for binding, washing and elution of the sample. RNA elution step was performed twice with 10 µL of nuclease free water, allowing 5 minutes of incubation time per elution to maximize recovery.

**Microfluidic chip fabrication for nanopore measurements**

A polydimethylsiloxane (PDMS) microfluidic chip was fabricated for nanopore measurements. The chip contained small chambers arranged in a radial geometry with respect to a central chamber, connected to the rest through small channels. The chip was produced by combining a 10:1 volume ratio of PDMS monomer and curing agent (Sylgard 184 silicone elastomer kit, The Dow Chemical Company), which were mixed for 10 minutes and then poured into a preheated mould (60 °C, 3 minutes). The mould was put into a vacuum to remove air bubbles in PDMS and then it was heated at 60 °C for 48 hours. The PDMS chips were removed from the mould and small perpendicular holes were created in the channels connecting the chambers. Then, glass nanopores were placed in the channels and the chip was sealed by pressing it against a glass slide after being treated for 11 seconds in a plasma chamber using maximum power (Femto, Diener). The channels were then sealed by introducing the PDMS mixture (10:1) in the perpendicular orifices and the chip was heated in a hot plate at 150°C for 5 minutes, followed by 100°C heating for 60 minutes. After cooling down, the chip was treated for 5 min in plasma chamber at maximum power and all the chambers were filled with 4 M LiCl.

**Agarose gel electrophoresis**

RNA, DNA, and RNA ID samples were run on a 1% (w/v) agarose gel prepared in fresh 1 × TBE buffer, with 0.02 % sodium hypochlorite using autoclaved Milli-Q water for both the gel and running buffer preparation. The samples were run with 1 × TriTrack loading dye (Thermo Fisher, Catalog number R1161). A constant voltage of 70 V was applied for 180 minutes, and 150 ng of the sample was added to each lane. The gel was stained in 3 × GelRed buffer (Biotium) and imaged with a GelDoc-It™(UVP). Gel images were processed using ImageJ

(Fiji)[1]. The grayscale was inverted, the contrast was increased, the background was subtracted with a rolling ball of 50-150 and the surface was smoothened.

**Single-molecule sizing**

A python-based graphic user interface was used for single-molecule sizing of RNA ID. In each translocation event, downward current spikes associated to components of RNA ID design with known base pair distance were selected manually. The known base pair separation between the spikes was associated with the time distance between both spikes, to obtain a base pair-to-time conversion factor. The start and end points of each event were selected manually, and the total event translocation time was converted to an estimate of base pairs using the respective conversion factor for each RNA ID translocation.

**Figure 1.**

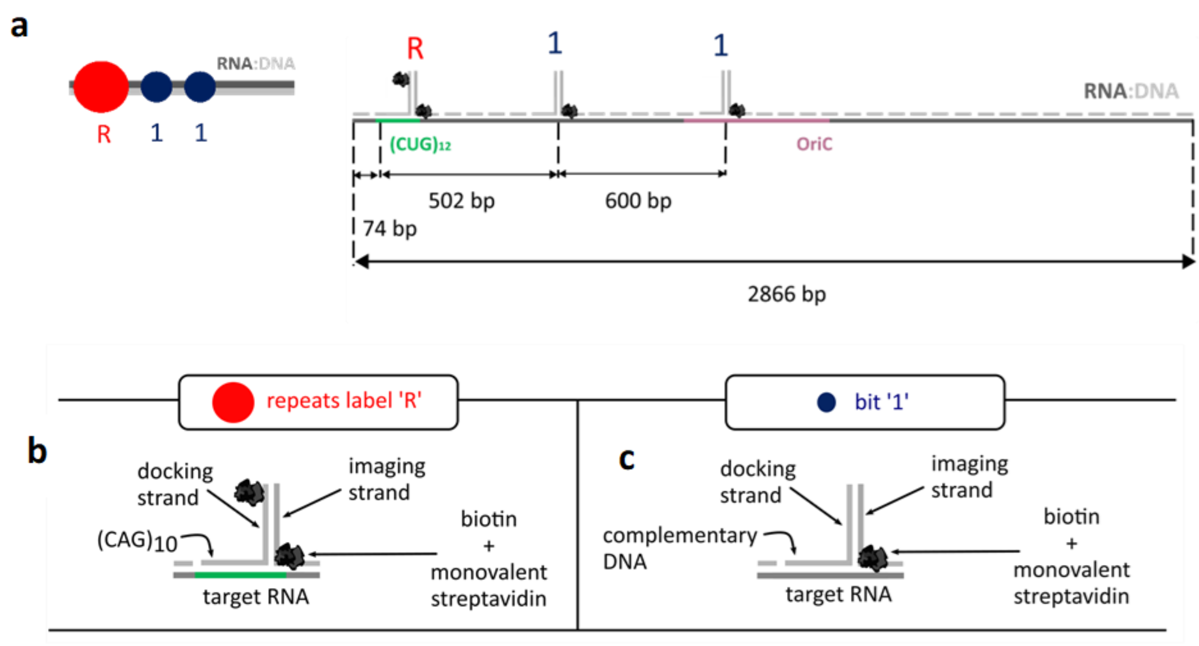


**Figure 1.** Design of RNA ID. **a** The position of the repeats label ‘R’ and the ‘1’ bits within the RNA ID is shown. **b** ‘R’ represents a DNA $CAG_{10}$ oligonucleotide (docking strand) that binds to the CUG repeats, this oligo has an overhang sequence with 3’ biotin that binds to a complementary strand (imaging strand) with another 3’ biotin, enabling the overall binding of two monovalent streptavidins to RNA ID. **c** The RNA ID is decorated with ‘1’. Two ‘1’ bits are included in the RNA ID design to produce a distinguishable signal. ‘1’ bits lack biotin on the docking strand, which enables the binding of only one monovalent streptavidin per bit, providing a discriminatory signal in nanopore recordings.

#### Figure 2.

**a**

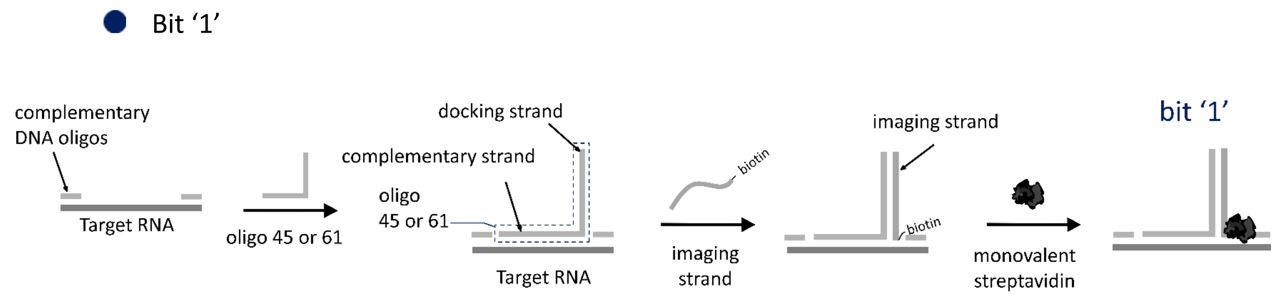


**b**

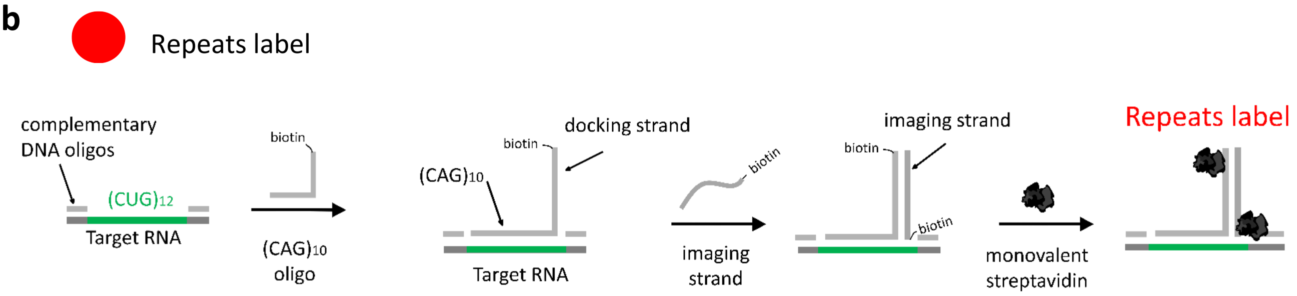


**c**

| **Oligo name** | **Sequence (5'→ 3')** |
|---|---|
| Docking strand | TTTGGATATCACTCATTAGTGGT |
| Imaging strand | ACCACTAATGAGTGATATCC/3'-biotin/ |

**Figure 2.** Assembly of RNA ID. **a** Detailed assembly of '1' bits. The sequence of each '1' bit can be found in Supplementary Table 2, corresponding to oligos 45 and 61. **b** Detailed assembly of repeats label 'R', sequence can be found in Supplementary Table 2 (oligo 76). **c** Table shows the sequence of docking strand overhang and its complementary oligo (imaging strand) which enable streptavidin binding.

**Figure 3.**

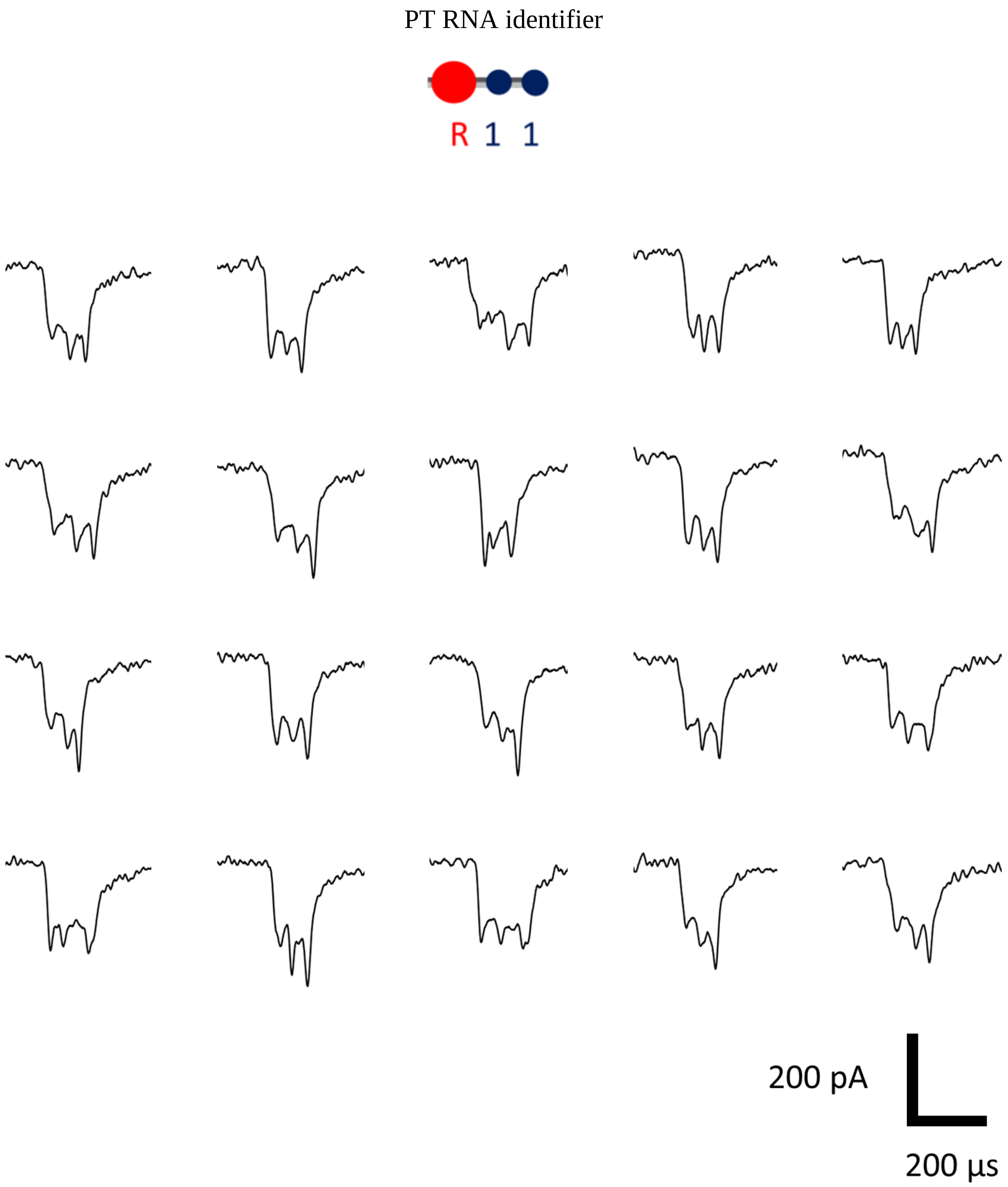


**Figure 3.** Nanopore translocation events of RNA identifier from premature transcription termination (PT). These are the first 20 unfolded translocation events detected of PT RNA IDs. Nanopore event variability can be accounted to the physical configuration-dependent RNA ID transport[2].

**Figure 4.**

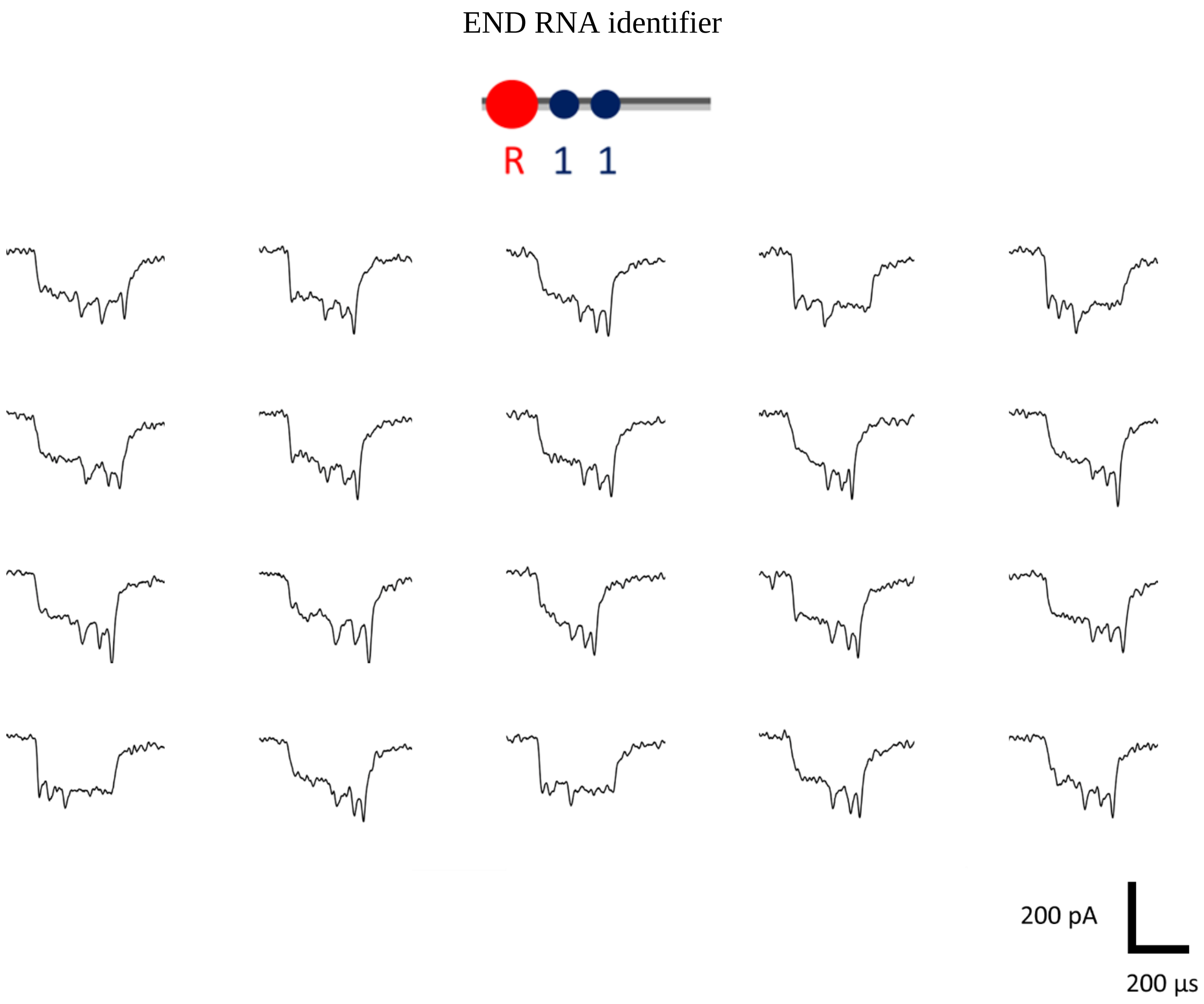


**Figure 4.** Nanopore translocation event of RNA identifier from transcription of the complete linear DNA (END). T7RNAP falls off at the end of the linear template causing transcription termination. These are the first 20 unfolded translocation events detected of END RNA IDs.

**Figure 5.**

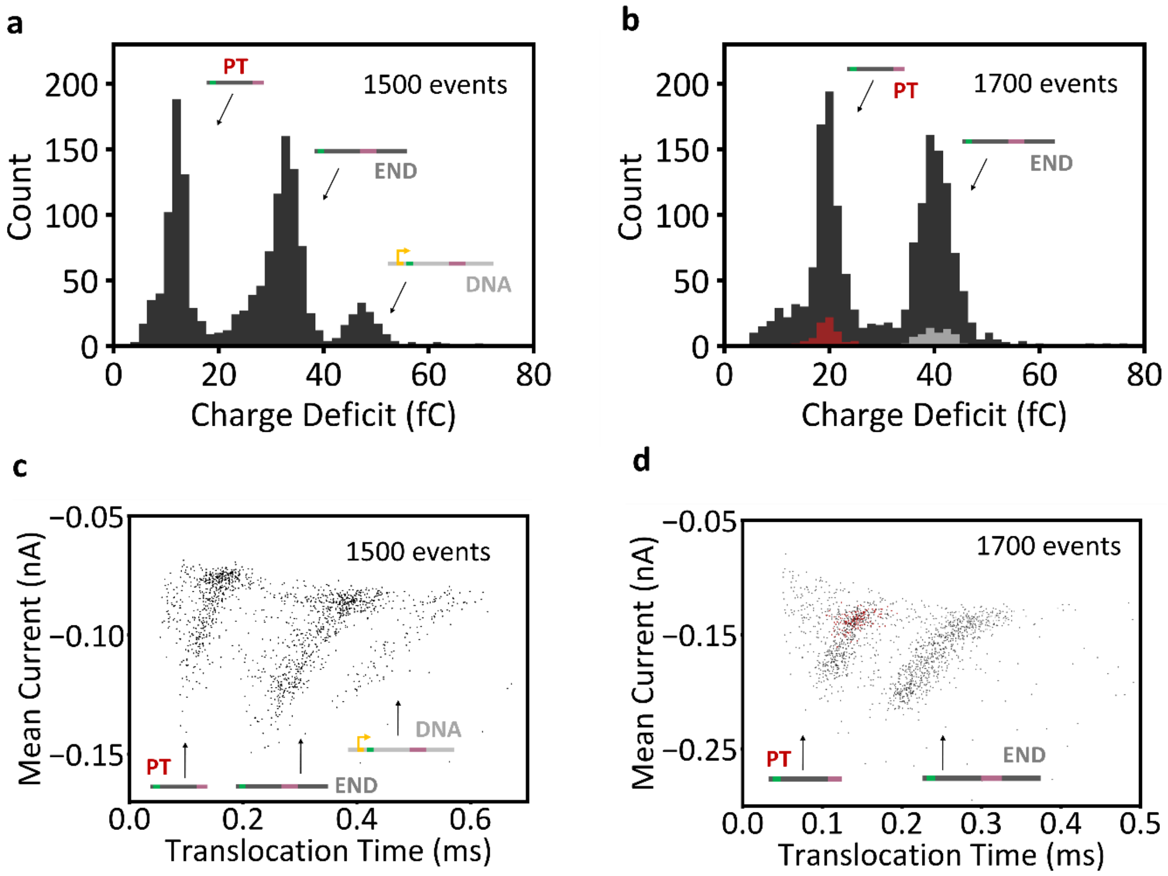


**Figure 5.** Characterization of RNA ID using charge deficit, mean current and translocation time. **a** Histogram of the charge deficit of RNA ID still in the presence of the linear DNA template. Histogram shows 3 distributions, the one with the lowest charge deficit is ascribed to premature termination (PT), the middle distribution corresponds to transcription of the full linear DNA (END) and the distribution furthest to the right is ascribed to the linear DNA template. This distribution (composed of 1500 events) includes translocations of molecules with multiple conformations, which include folded events, constructs with knots and unfolded events. **b** After treatment with DNase I, it can be seen how the distribution furthest to the right, ascribed to DNA, is removed. Unfolded translocations of both PT (red) and END (gray) RNA IDs (presented in Figure 2c) describe the entire sample, despite their conformation, while enabling single-molecule sizing. These distributions correspond to the same nanopore measurement presented in Figure 2. **c** Scatter plot of mean current against translocation time shows three distinct distributions, attributed to PT RNA IDs, END RNA IDs, and the linear template (from left to right). Unfolded molecules take longer to translocate through the pore than folded molecules but cover less cross-sectional area of the pore while translocating,

producing a less significant drop in ionic current for longer times. Plotting events with different conformations produces this type of distribution, with unfolded events at the top, and folded events at the bottom of each distribution. **d** Selection of unfolded events (in the sample treated with DNase I) is performed to describe each distribution. Unfolded events were found at the top of each distribution, and they are representative of the whole sample.

**Figure 6.**

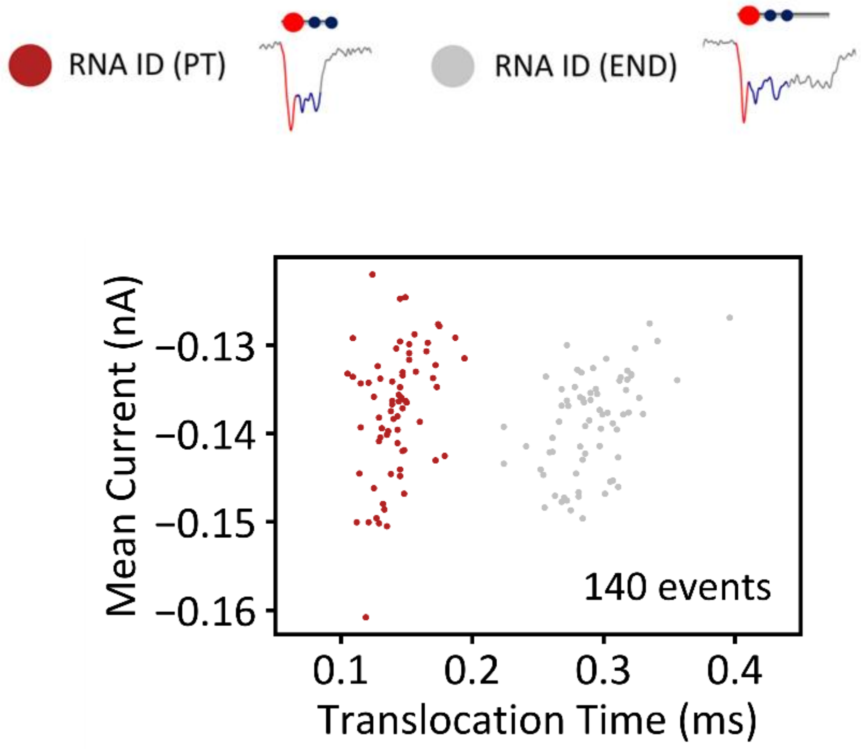


**Figure 6.** Scatter plot of mean current against translocation time for unfolded END (red) and PT (gray) RNA IDs, which depicts two distinct populations. This demonstrates that plotting these two parameters for transcripts with different termination sites enables their distinction.

#### Figure 7.

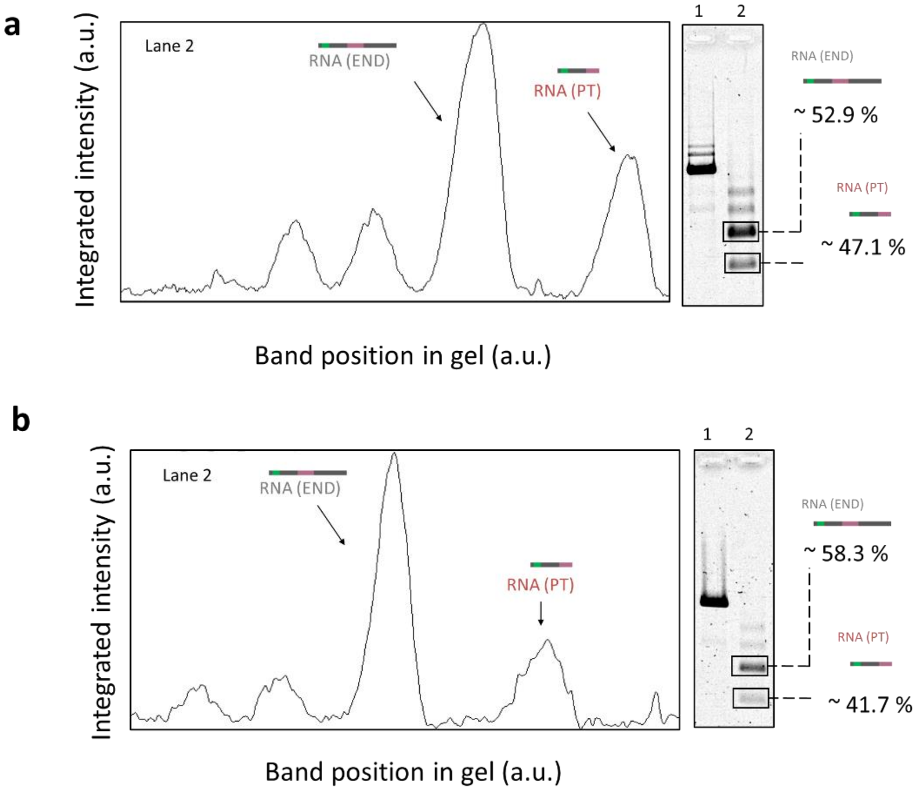


For **a**

| Lane | Molecule | Intensity profile (a.u.) | Normalized intensity profile (a.u./bp) |
|---|---|---|---|
| 1 | RNA (PT) | 8181.711 | 5.389 |
| | RNA (END) | 17336.317 | 6.049 |

For **b**

| Lane | Molecule | Intensity profile (a.u.) | Normalized intensity profile (a.u./bp) |
|---|---|---|---|
| 1 | RNA (PT) | 5107.296 | 3.364 |
| | RNA (END) | 13481.004 | 4.703 |

**Figure 7.** Quantitative analysis of premature transcription termination in OriC using agarose gel electrophoresis. **a** The gel shows the linearized construct in lane 1 and the DNase I treated transcripts in lane 2. The two topmost bands in lane 2 are attributed to RNA side products. The two lower bands correspond to END RNA and PT RNA, from top to bottom. The intensity

profile of these two bands was plotted and the area of each peak was computed. The peak areas were normalized by the number of base pairs of each transcript to obtain an estimate of transcript abundance. Gel suggests premature transcription termination of ~ 47.1 % in OriC. The same experimental procedure and analysis described was repeated to evaluate variability in RNA abundance quantification. Gel suggests premature transcription termination of ~ 41.7 % in OriC, indicating good reproducibility. Lane 1 – linear DNA, Lane 2 - DNase treated transcripts. Gel: 1 % (w/v) agarose, 1 × TBE, 0.02% sodium hypochlorite.

#### Figure 8.

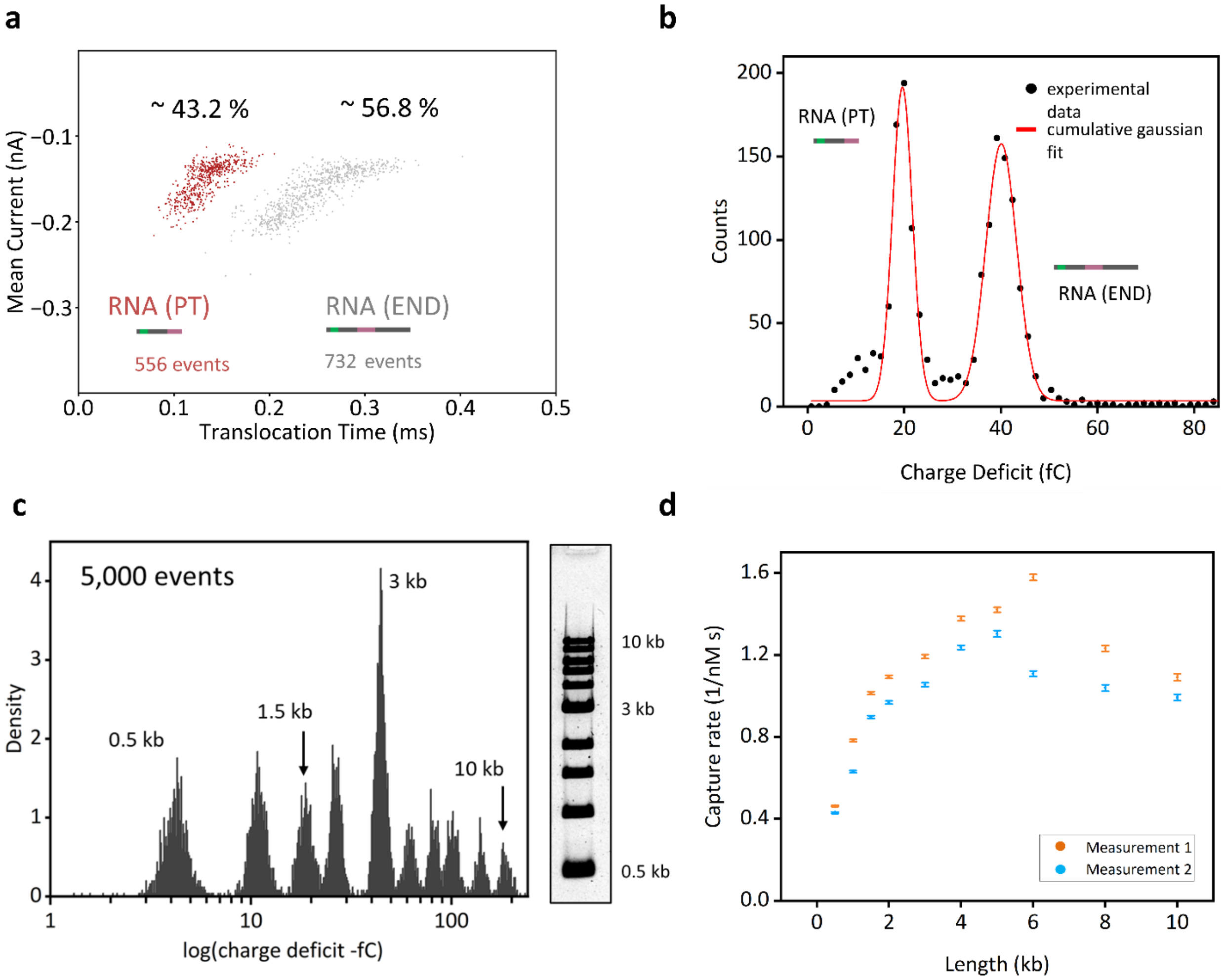


**Figure 8.** Quantitative study of PT RNA IDs and END RNA IDs events detected in nanopore sensors. **a** As illustrated in Supplementary Figure 5c and Supplementary Figure 5d, scatter plots of mean current against translocation time can be used to identify RNA IDs from transcripts with different termination sites. Scatter plot exhibits two distributions: the one from the left is ascribed to PT RNA ID and the distribution at the right corresponds to END RNA ID. The events within each of these distributions were counted by establishing lower and upper limits in charge deficit, the rest of the events were discarded. The upper limit for RT RNA IDs was 25 fC, and the lower limit was 15 fC. For END RNA IDs the upper limit was 50 fC and the lower limit was 30 fC. 556 events were detected for RT RNA IDs and 732 events were identified for END RNA IDs, suggesting ~ 43.2% termination in OriC. **b** Cumulative Gaussian fit of PT and END RNA IDs. The lowest charge deficit peak is ascribed to premature termination (PT) and the middle peak corresponds to full-length RNA transcript (END). Comparison of the areas of PT RNA ID and END RNA ID fits suggest ~44% transcription

termination. **c** A 1kb DNA ladder (NEB) containing DNA molecules from 0.5 kb to 10 kb was used to study capture rate variability in our nanopore system ascribed to a difference in length-dependent capture rate[3,4]. The concentration is known for the DNA of each length, based on information provided by the manufacturer. Characterization of the DNA ladder in nanopores was performed in duplicate. **d** Capture rate for each DNA molecule length included in the DNA ladder for both measurements.

**Figure 9.**

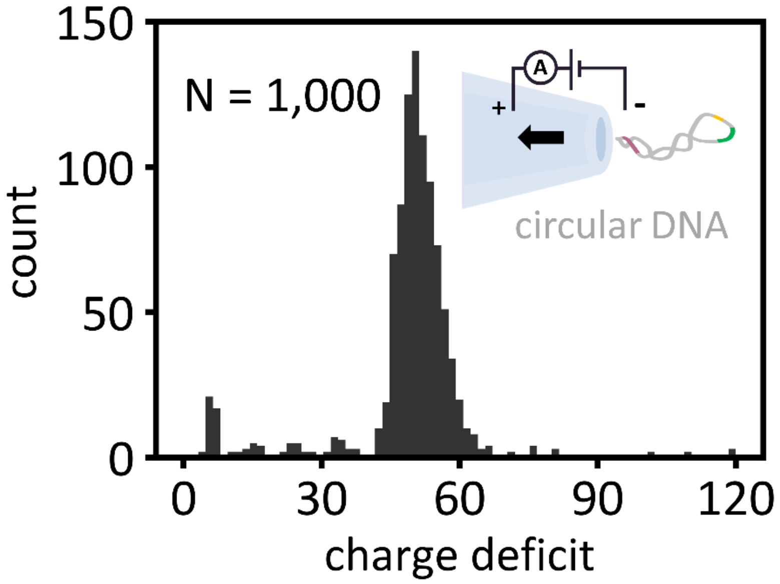


**Figure 9.** The 3.1 kbp circular DNA used as the template for rolling circle transcription was characterized using nanopore sensing. The charge deficit of the translocation events shows a unimodal distribution, demonstrating the presence of a single DNA construct. This also confirms RNA is produced from circle rolling transcription of the 3.1 kbp circular DNA, and not by transcription of DNA dimers or trimers of the 3.1 kbp DNA which could be a possible interpretation of agarose gel electrophoresis. Here we also showcase the clarity nanopore sensing provides for the elucidation of DNA identity.

**Figure 10.**

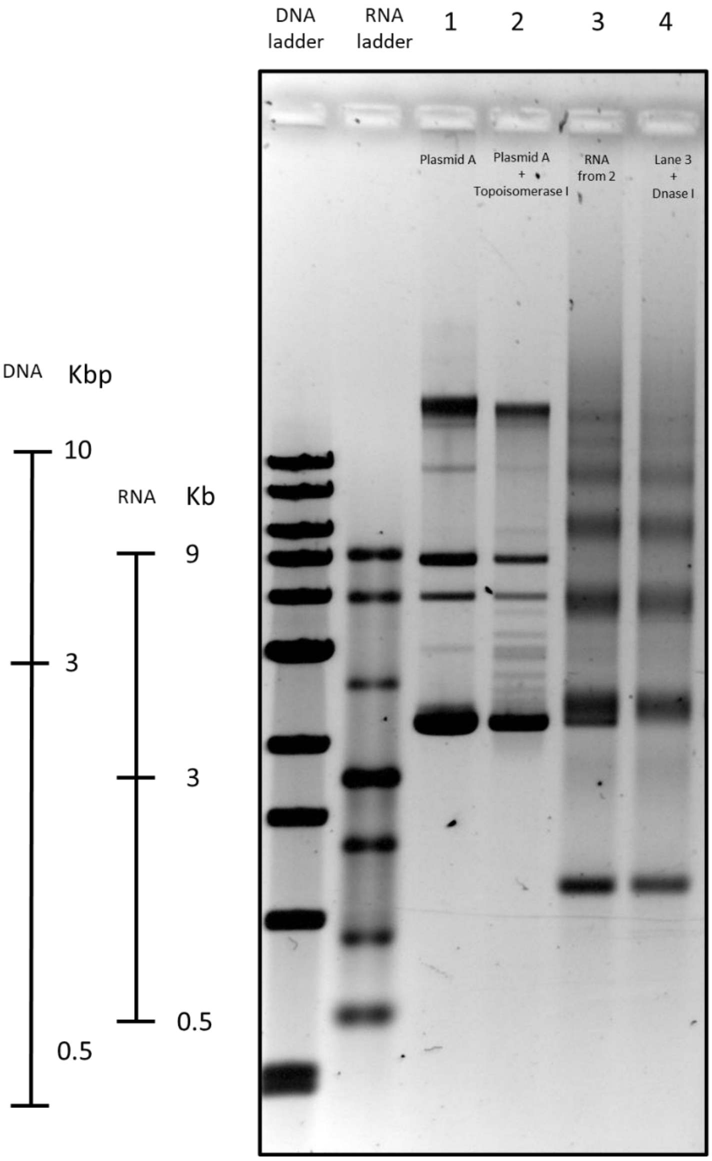


**Figure 10.** Rolling circle transcription of circular DNA construct. DNA ladder and ssRNA ladder are included on both sides of the gel. Lane 1 – Circular plasmid with 12 CTG repeats (sequence of circular plasmid in Supplementary Table 1). The multiple bands are ascribed to the physical configurations that supercoiled DNA has while being electrophoretically driven through the agarose gel. Lane 2 – circular plasmid from lane 1 treated with Escherichia coli Topoisomerase I to induce plasmid relaxation. Lane 3 – RNA from transcription of Topoisomerase I treated circular plasmid in lane 2. Lane 4 – RNA from lane 3 treated with DNase I. From DNase I treatment, DNA band located at ~3 kbp (DNA) is removed. The rest of the bands, ascribed to RNA products of the multiple transcription cycles, remain.

**Figure 11.**

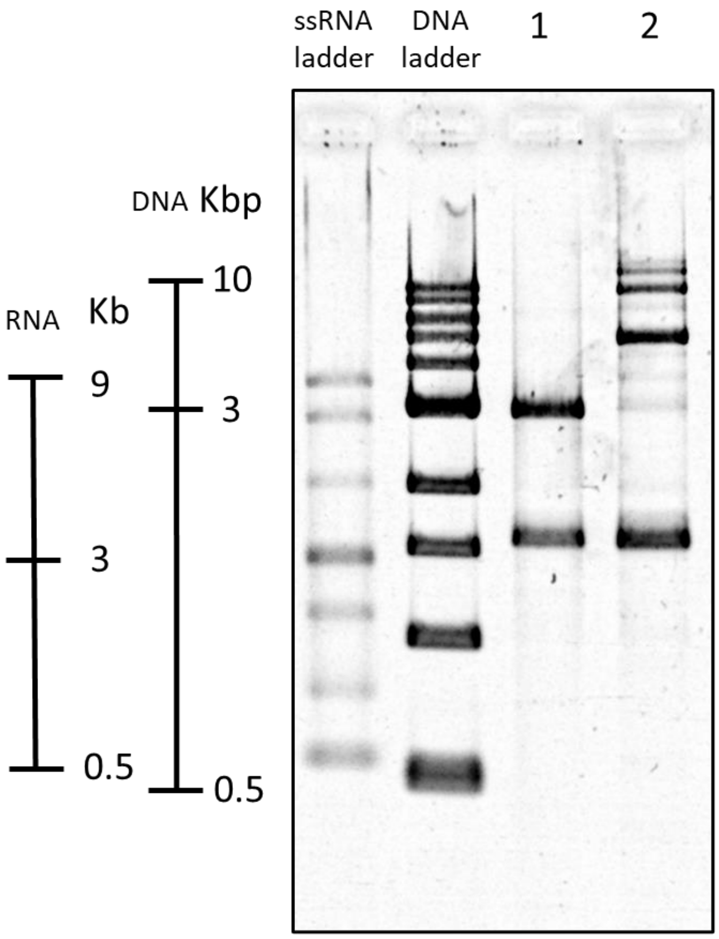


**Figure 11.** RNA ID of transcripts from transcription of linear DNA construct and rolling circle transcription of circular DNA construct (sequence in Supplementary Table 1). The agarose gel shows a DNA ladder and ssRNA and DNA ladder on the left side. Lane 1 – RNA ID assembled from RNA from transcription of linear DNA template. Lane 2 - RNA ID assembled from transcripts of rolling circle transcription. Gel: 1 % (w/v) agarose, 1 × TBE, 0.02% sodium hypochlorite.

**Figure 12.**

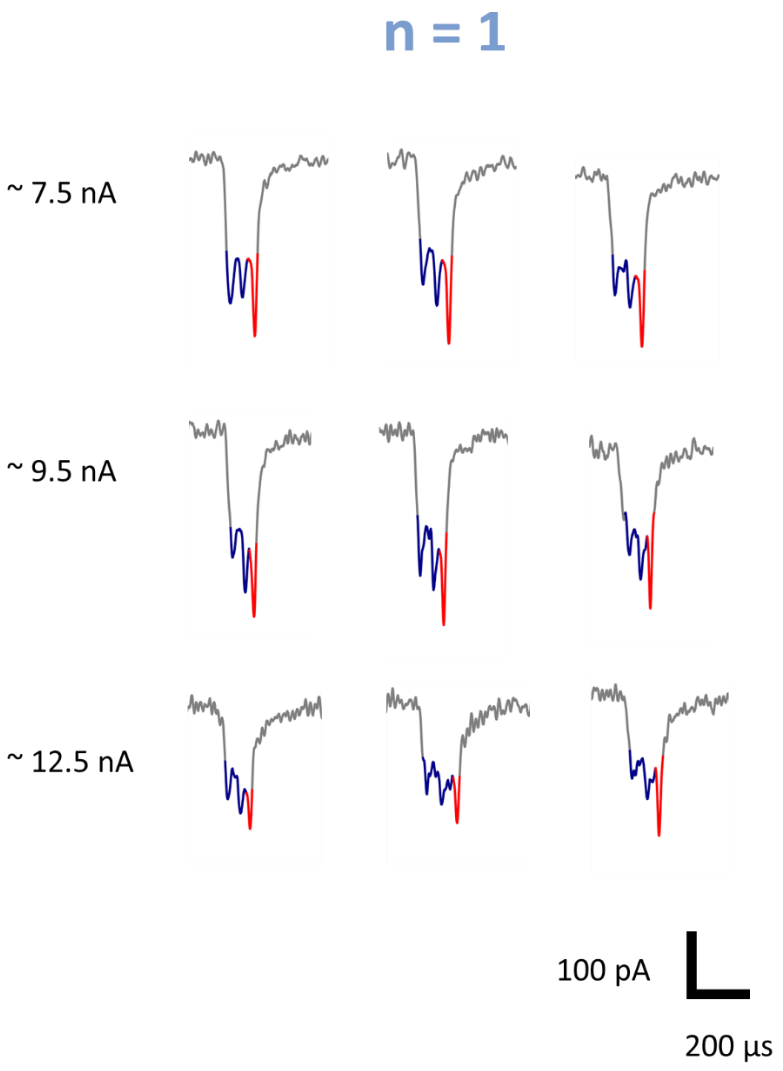


**Figure 12.** N = 1 example events. Example nanopore events of RNA IDs produced from one transcription cycle (N = 1) measured in pores with different sizes, which exhibit different signal to noise ratio. Variation in pore size can be inferred from changes in the ionic current baseline. All measurements were performed under the same applied voltage of 600 mV. Larger pores have a larger ionic current baseline.

**Figure 13.**

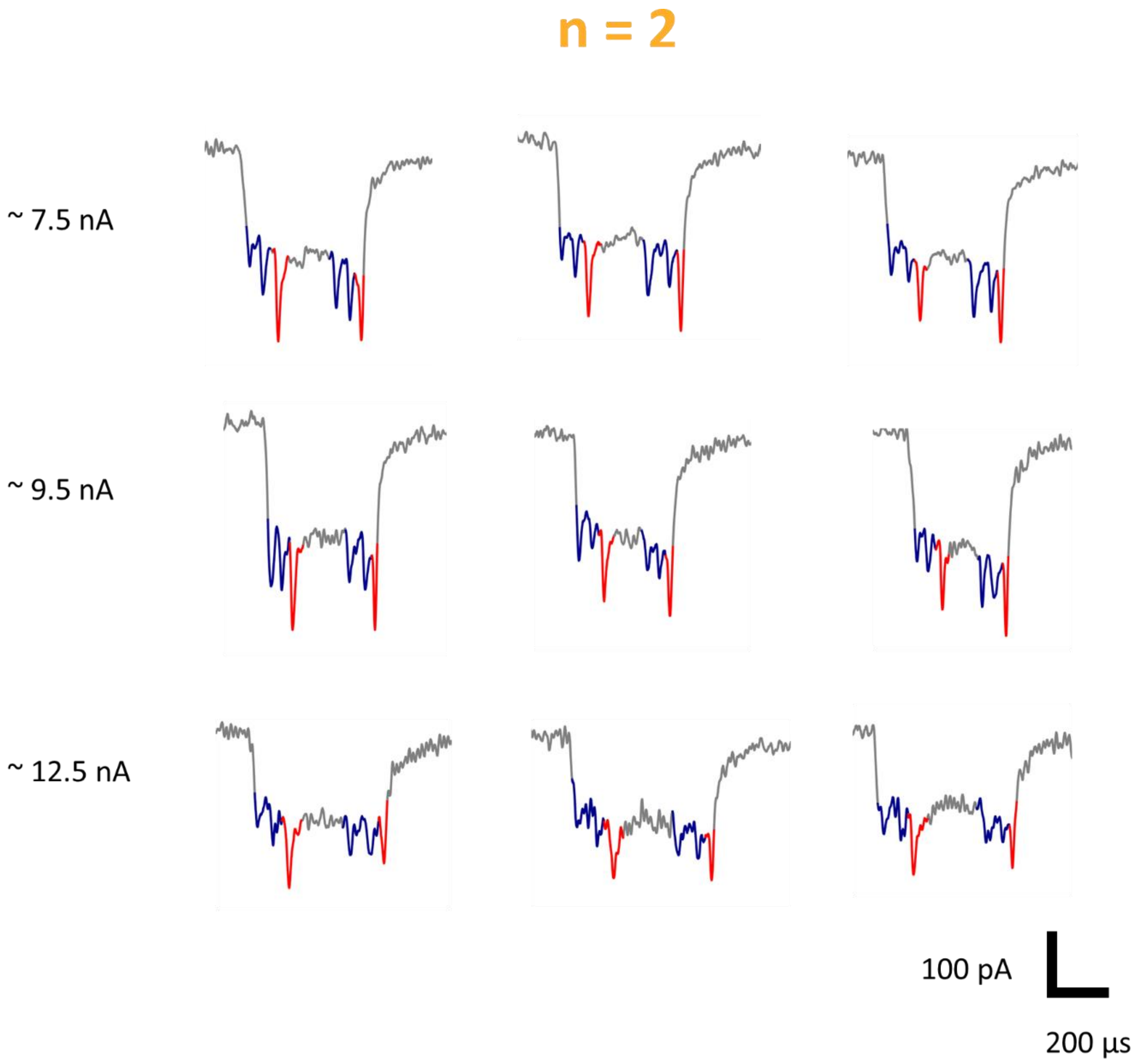


**Figure 13.** N = 2 example events. Example nanopore events of RNA IDs produced from two transcription cycles (N = 2) measured in pores with different sizes, which exhibit different signal to noise ratio. Variation in pore size can be inferred from changes in the ionic current baseline. All measurements were performed under the same applied voltage of 600 mV. Larger pores have a larger ionic current baseline.

**Figure 14.**

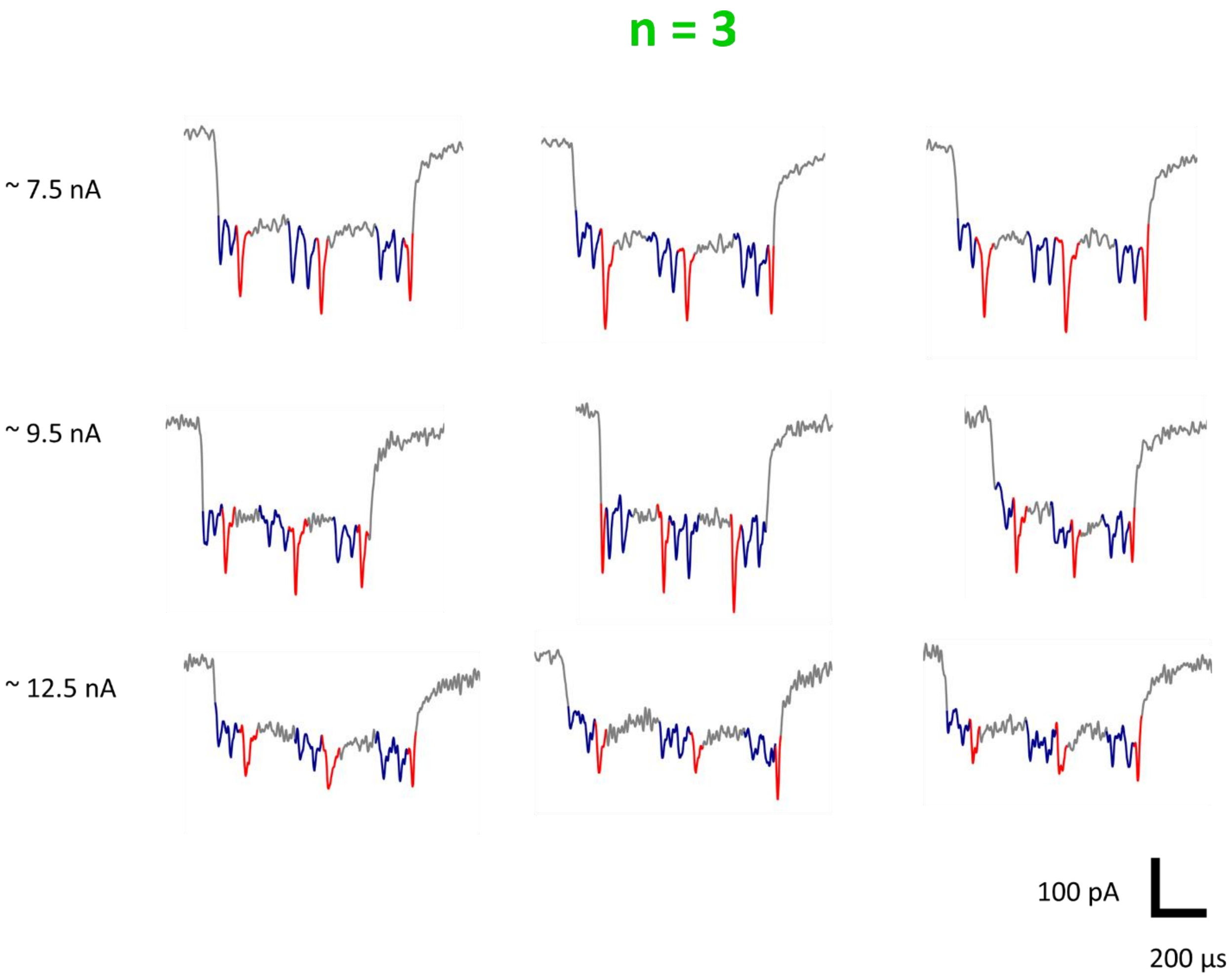


**Figure 14.** N = 3 example events. Example nanopore events of RNA IDs produced from three transcription cycles (N = 3) measured in pores with different sizes, which exhibit different signal to noise ratio. Variation in pore size can be inferred from changes in the ionic current baseline. All measurements were performed under the same applied voltage of 600 mV. Larger pores have a larger ionic current baseline.

**Figure 15.**

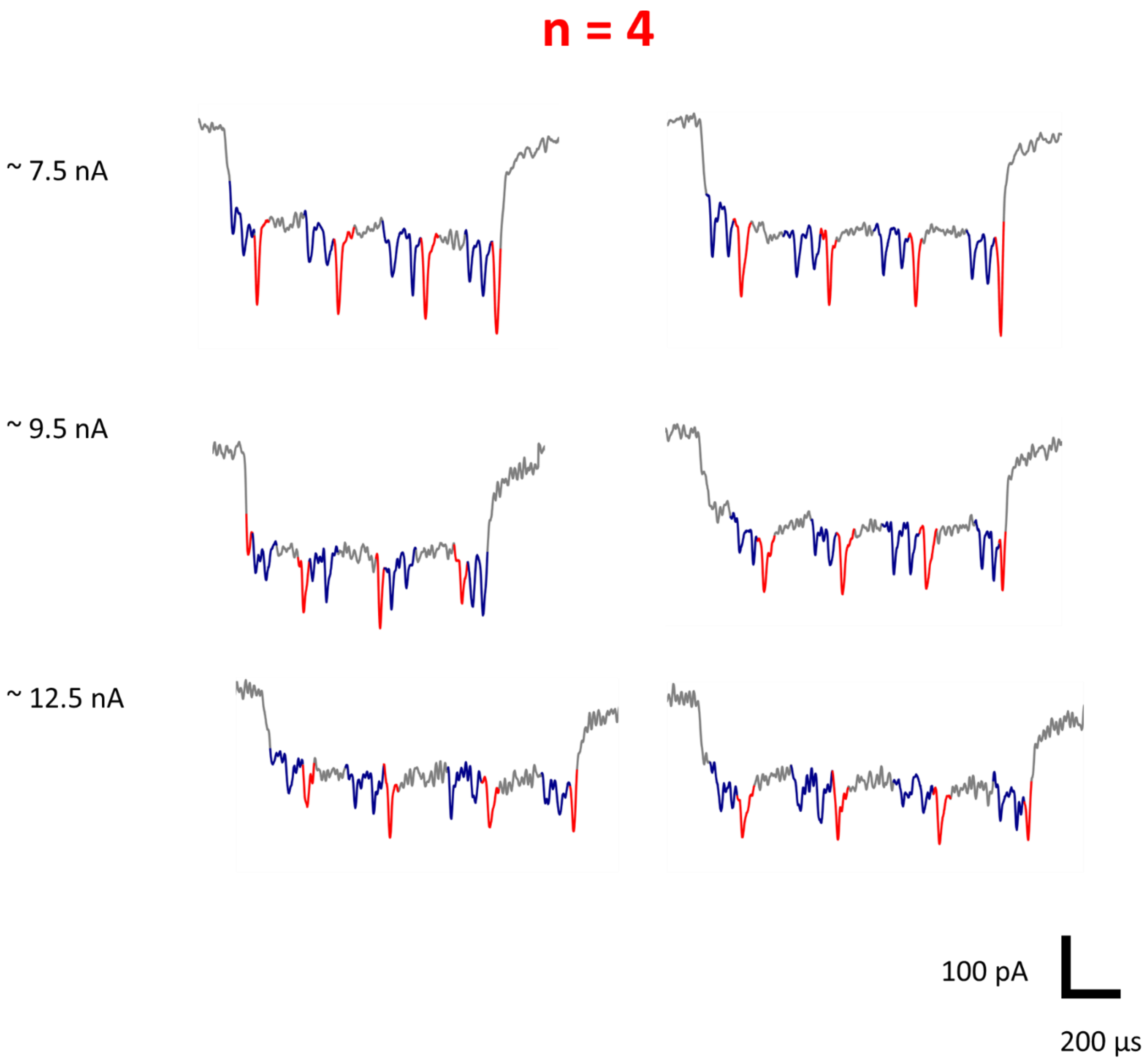


**Figure 15.** N = 4 example events. Example nanopore events of RNA IDs produced from four transcription cycles (N = 4) measured in pores with different sizes, which exhibit different signal to noise ratio. Variation in pore size can be inferred from changes in the ionic current baseline. All measurements were performed under the same applied voltage of 600 mV. Larger pores have a larger ionic current baseline.

**Figure 16.**

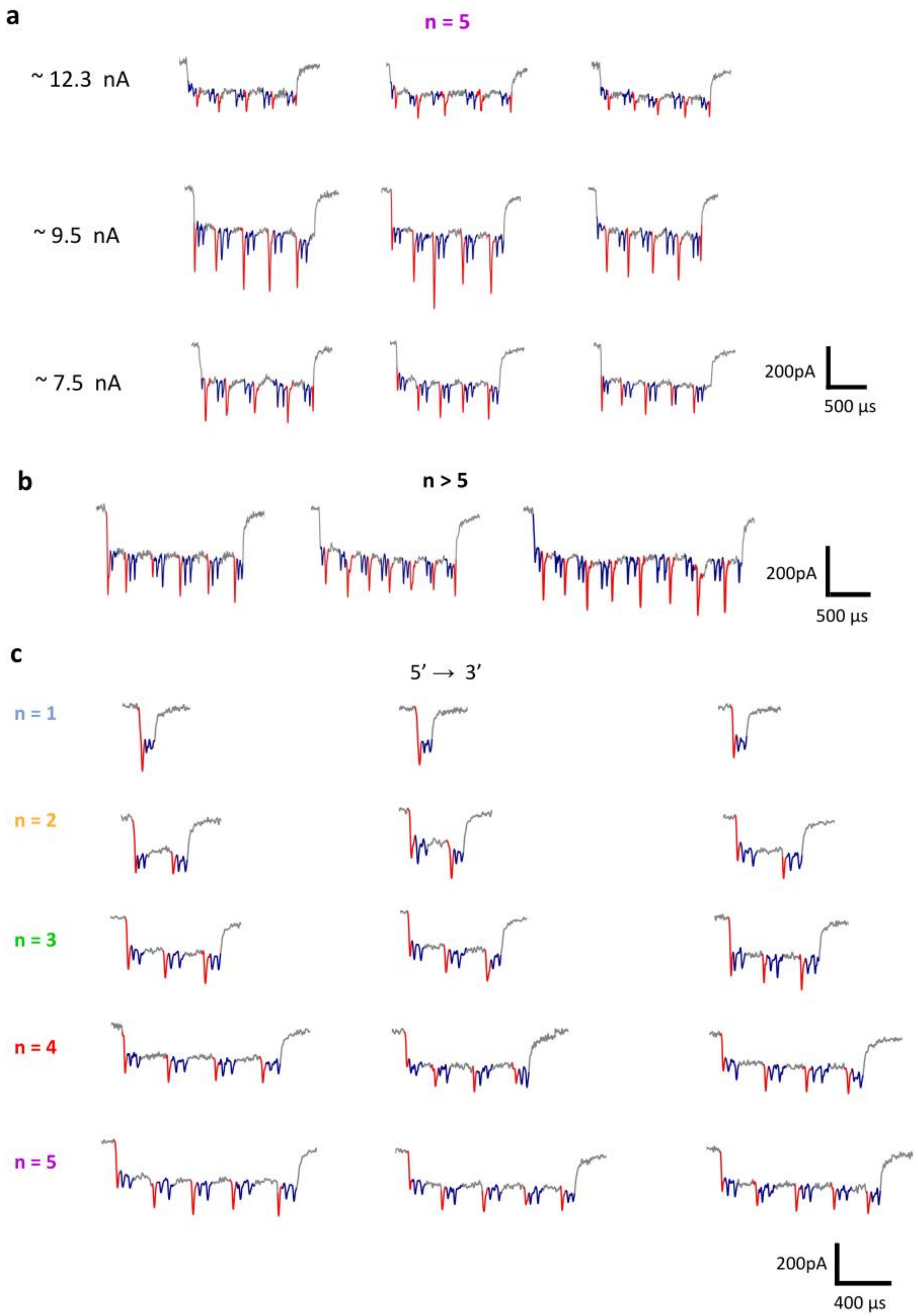
a
n = 5
~ 12.3 nA
~ 9.5 nA
~ 7.5 nA
200pA
500 µs
b
n > 5
200pA
500 µs
c
5′ ⟶ 3′
n = 1
n = 2
n = 3
n = 4
n = 5
200pA
400 µs

**Figure 16.** RNA ID nanopore events for transcription cycles N = 5 and N > 5 example events. **a** Example of nanopore events of RNA IDs produced from five transcription cycles (N = 5) measured in pores with different sizes. All measurements were performed under the same applied voltage of 600 mV. **b** Also, RNA IDs with N > 5 were identified, example events are shown from different nanopore measurements. **c** Translocation of RNA IDs for each transcription cycle that entered to the pore in the 5′ to 3′ direction.

**Figure 17.**

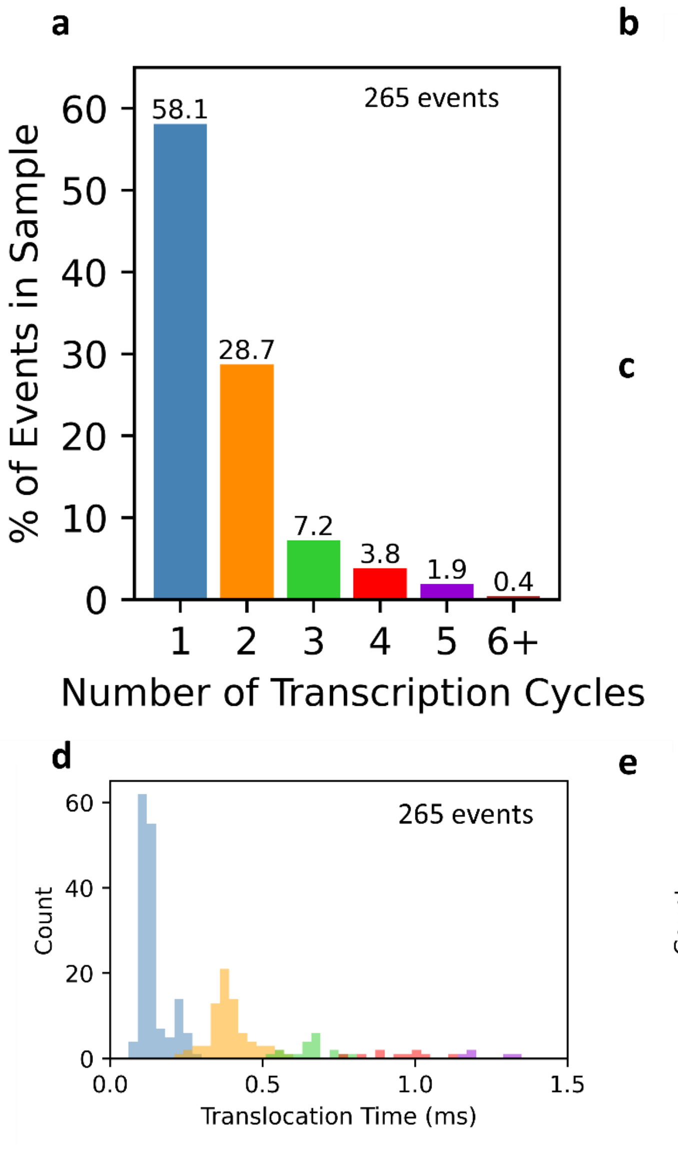


**Figure 17.** Translocation time and charge deficit of RNA IDs from rolling circle transcription. **a** For individual nanopore measurements presented in Figure 3e, 265 unfolded events were identified. Bar plots show the relative abundance of each RNA ID classified by number of transcriptions cycle. **b** Swarm plot of charge deficit per transcription cycle. **c** Swarm plot of translocation time per transcription cycle. **c** and **d** correspond to histograms of translocation time and charge deficit, respectively, presented in Figure 3e, but using a linear scale on the *y* axis instead of a logarithmic scale.

**Figure 18.**

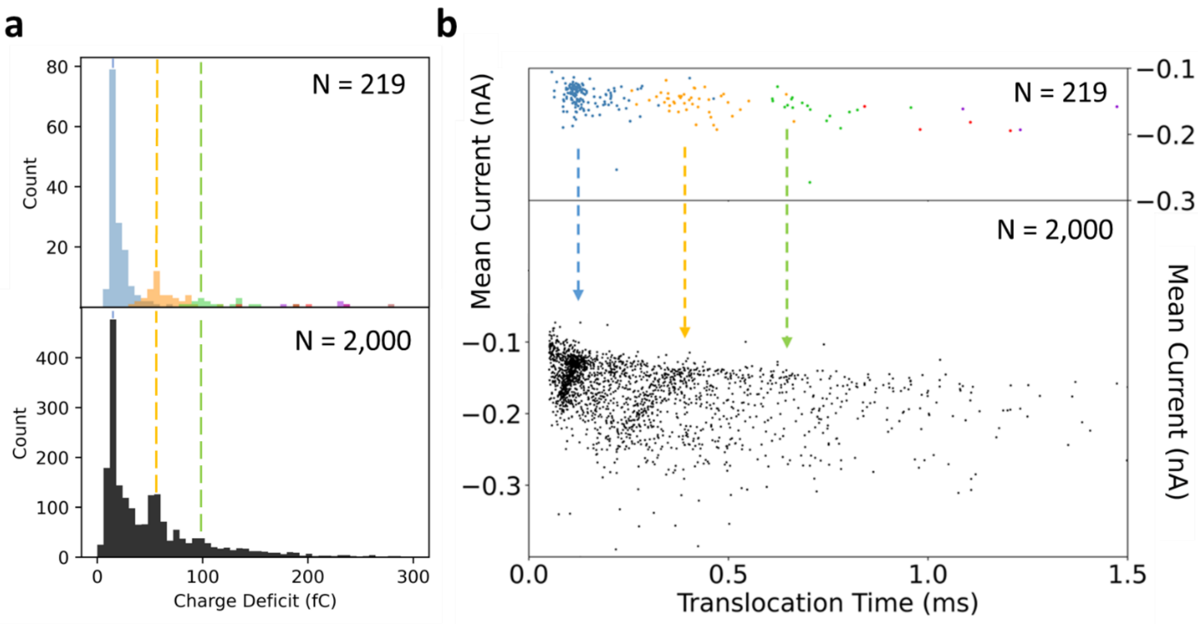


**Figure 18.** Unfolded (linear) RNA IDs events are representative of the events with different conformations. **a** Histogram of charge deficit for all translocations detected in one nanopore measurement (black, 2000 events). The selection of 219 unfolded events shows a distribution of charge deficit which is representative of the distribution of all translocations detected, therefore these events can be used to describe the sample and gain single-molecule information from the RNA ID design. The charge density distributions produced between the main distributions (green and yellow) are ascribed to fall-off of T7RNAP. **b** Scatter plot of mean current against translocation time, which also shows that the selection of unfolded can be used for the description of a sample within a defined parameter space.

**Figure 19.**

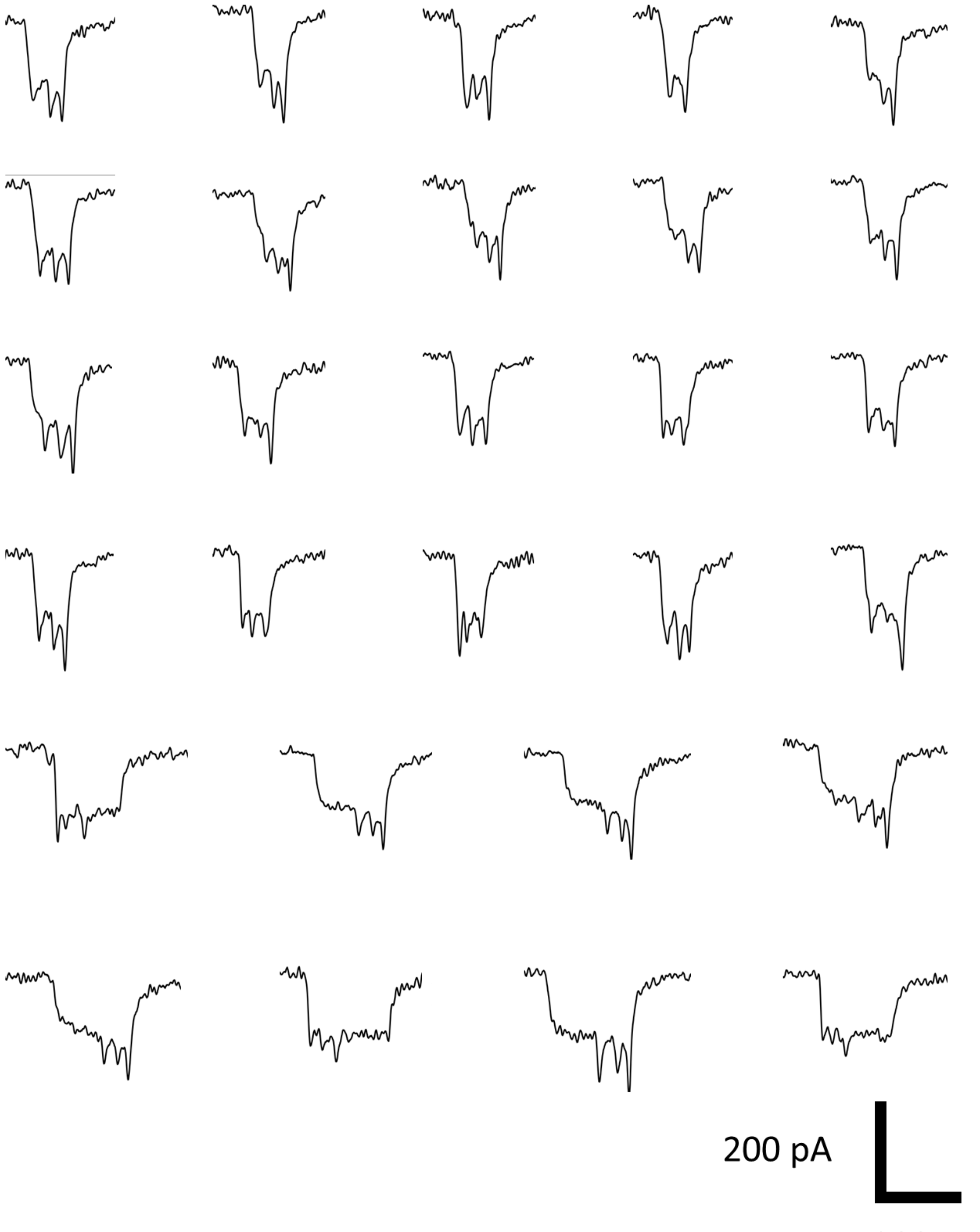


**Figure 19.** Example nanopore events of RNA IDs produced from one transcription cycle (N = 1) used for single-molecule sizing in Figure 4d. These are the first 28 unfolded translocation events detected for N = 1.

**Figure 20.**

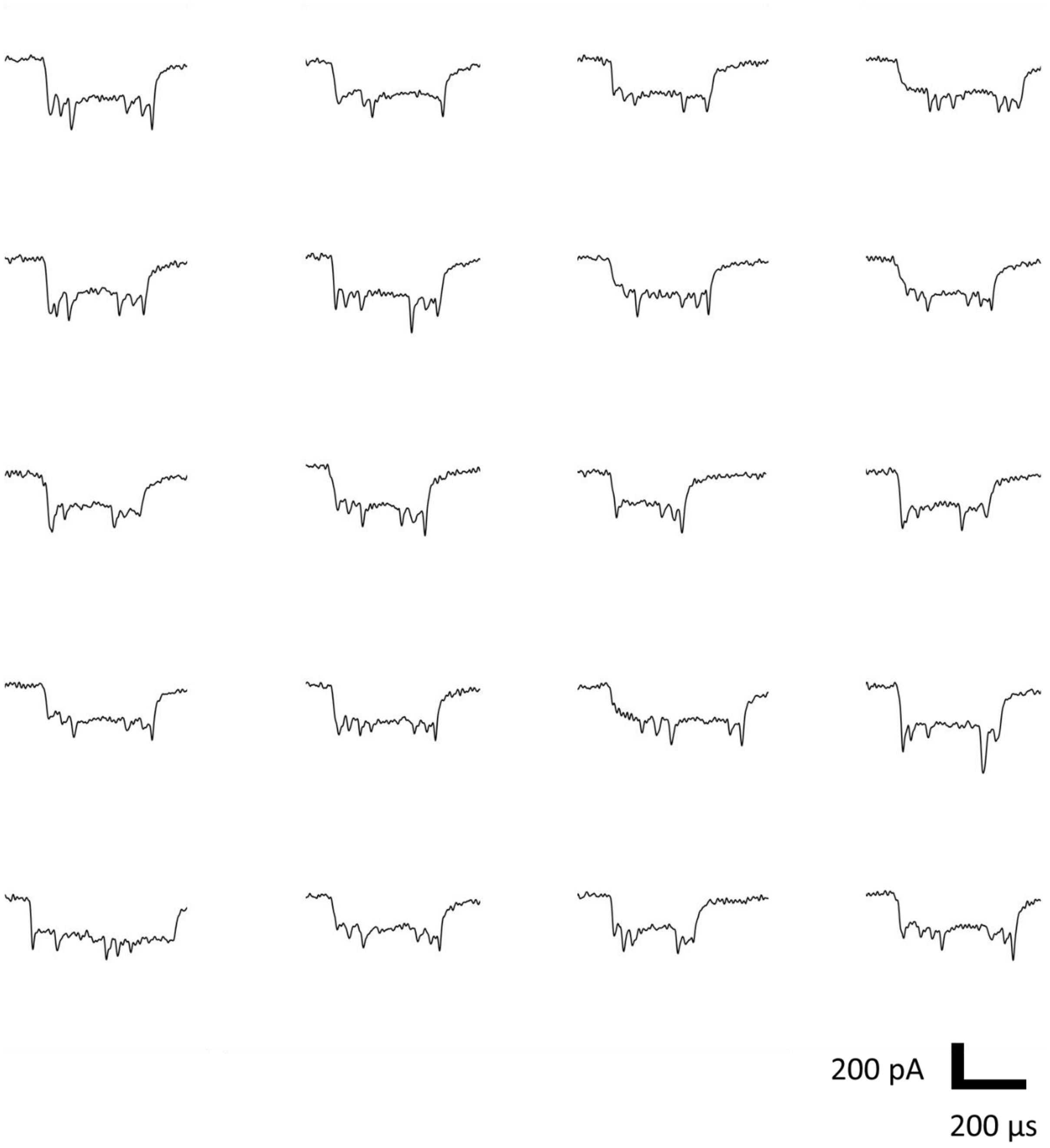


**Figure 20.** Example nanopore events of RNA IDs produced from two transcription cycles (N = 2) used for single-molecule sizing in Figure 4d. These are the first 20 unfolded translocation events detected for RNA IDs with N = 2.

**Figure 21.**

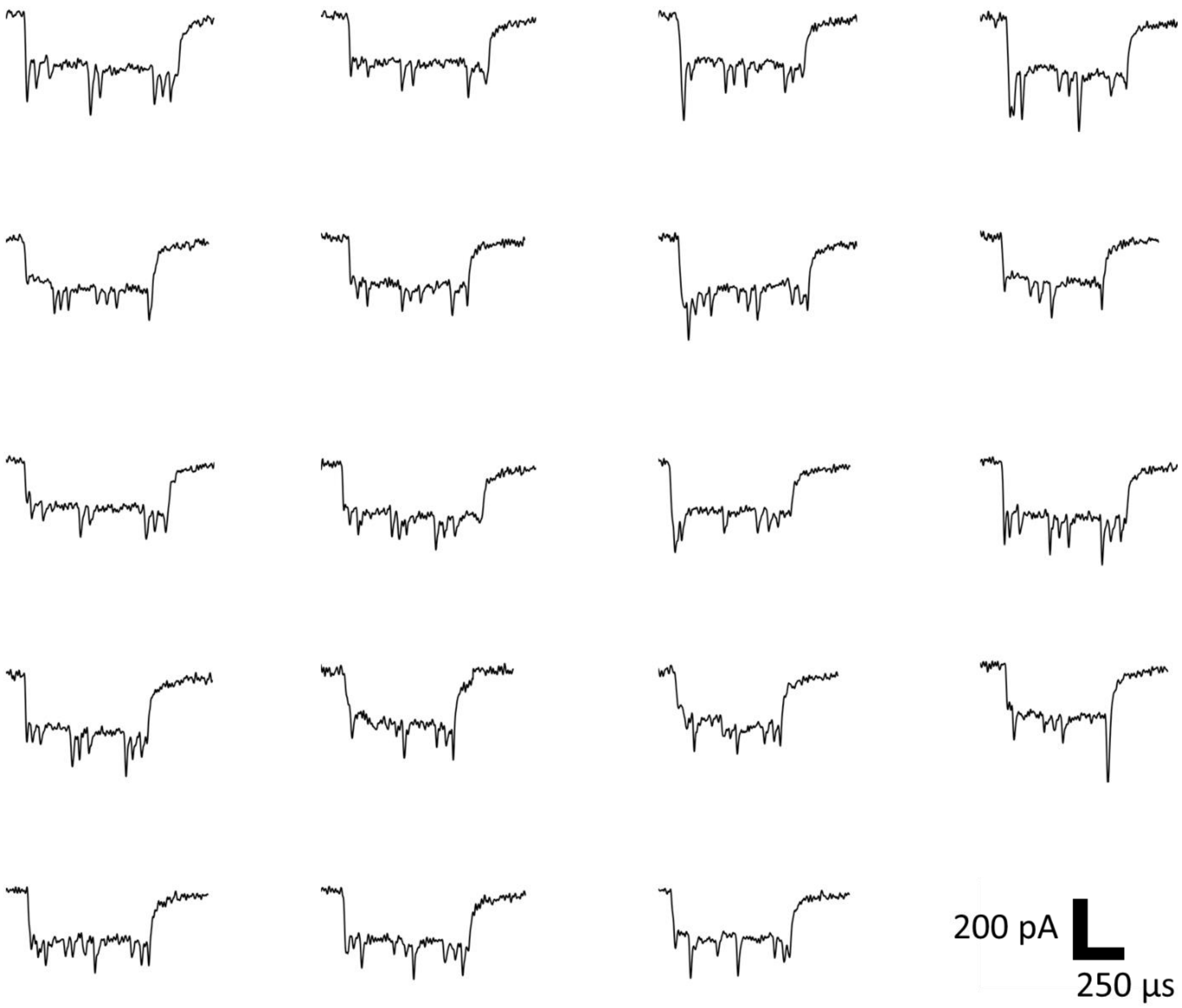


**Figure 21.** Example nanopore events of RNA IDs produced from three transcription cycles (N = 3) used for single-molecule sizing in Figure 4d. These are the first 20 unfolded translocation events detected for RNA IDs with N = 3.

**Figure 22.**

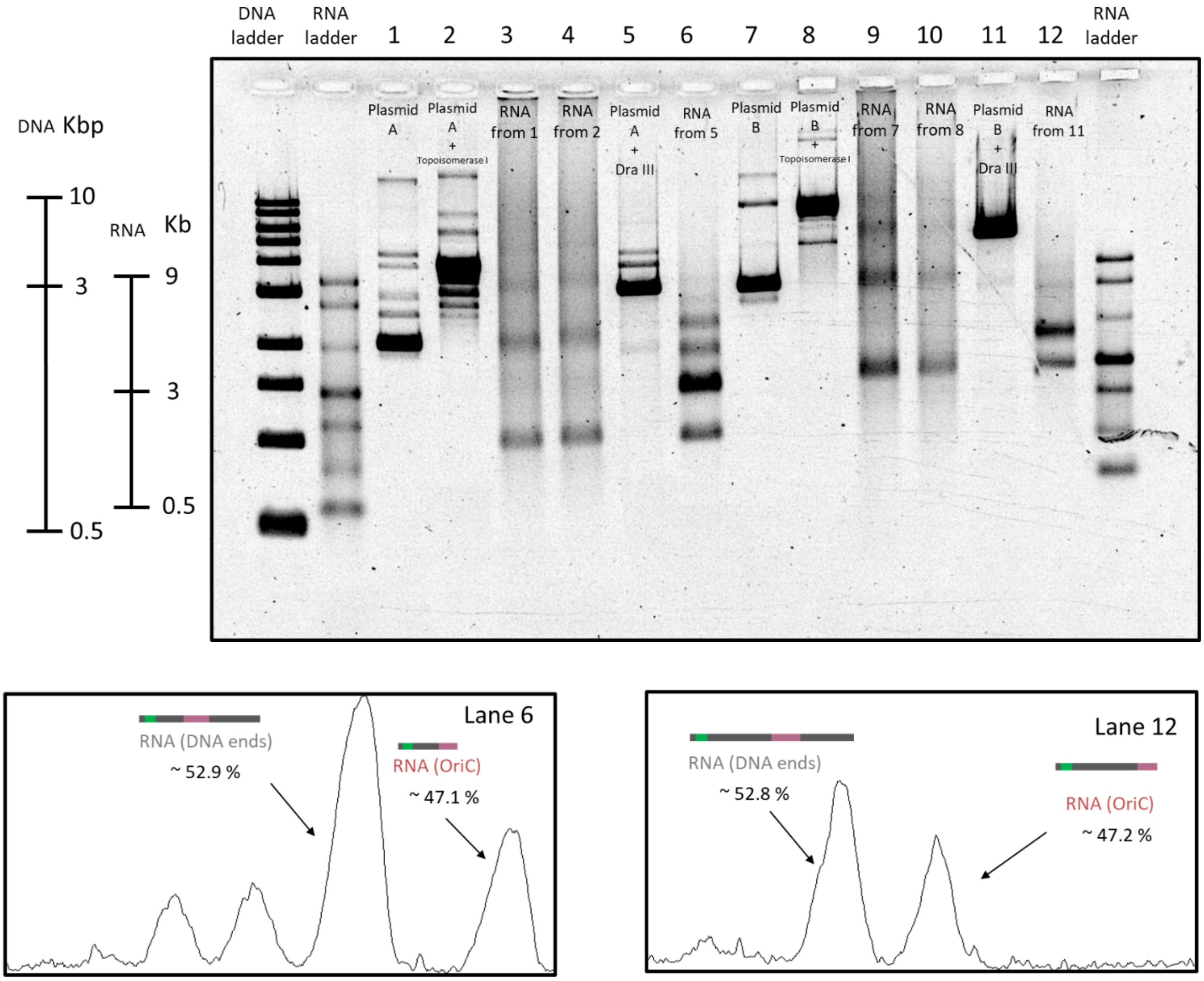


| Plasmid | Molecule | Intensity profile (a.u.) | Normalized intensity profile (a.u./bp) |
|---|---|---|---|
| A | RNA (OriC) | 8181.711 | 5.389 |
| | RNA (Template end) | 17336.317 | 6.049 |
| B | RNA (OriC) | 5889.418 | 1.983 |
| | RNA (Template end) | 9616.418 | 2.221 |

**Figure 22.** Transcription of circular and linear DNA templates with the OriC at different distances from the T7 RNA polymerase promoter. DNA ladder and ssRNA ladder are included. Lane 1 – circular DNA plasmid with 12 CTG repeats (sequence in Supplementary Table 1). Lane 2 – circular plasmid from lane 1 treated with *Escherichia coli* Topoisomerase I. Lane 3 – DNase I treated RNA from transcription of circular plasmid in lane 1. Lane 4 – DNase I treated RNA from transcription of Topoisomerase I treated circular plasmid in lane 2. Lane 5 – linear

DNA digested using DraIII-HF. Lane 6 – DNase treated RNA from transcription of linear DNA in lane 5. Lane 7 – 4527 kbp circular DNA plasmid (sequence in Supplementary Table 4). Lane 8 – circular plasmid from lane 7 treated with *Escherichia coli* Topoisomerase I. Lane 9 – DNase I treated RNA from transcription of circular plasmid in lane 7. Lane 10 – DNase I treated RNA from transcription of Topoisomerase I treated circular plasmid in lane 8. Lane 11 – linear DNA digested using DraIII-HF, it is a longer linear DNA than the linear template in lane 5. Lane 12 – DNase treated RNA from transcription of linear DNA in lane 11. The topmost band of lane 12, slightly below 5 kb, corresponds to transcription of the entire linear template, while the band from the bottom, allocated slightly below 3 kb, corresponds to premature termination in the OriC. The position of the bands is in good agreement with the expected premature termination in the OriC (see sequence in Supplementary Table 4). Gel: 1 % (w/v) agarose, 1 × TBE, 0.02% sodium hypochlorite. The intensity profile of both bands in lanes 6 and 12 were plotted and the area of each peak was computed. The peak areas were normalized by the number of base pairs of each transcript to obtain an estimate of transcript abundance. Gel suggests premature transcription termination of ~ 47 % in the OriC in both cases.

**Figure 23.**

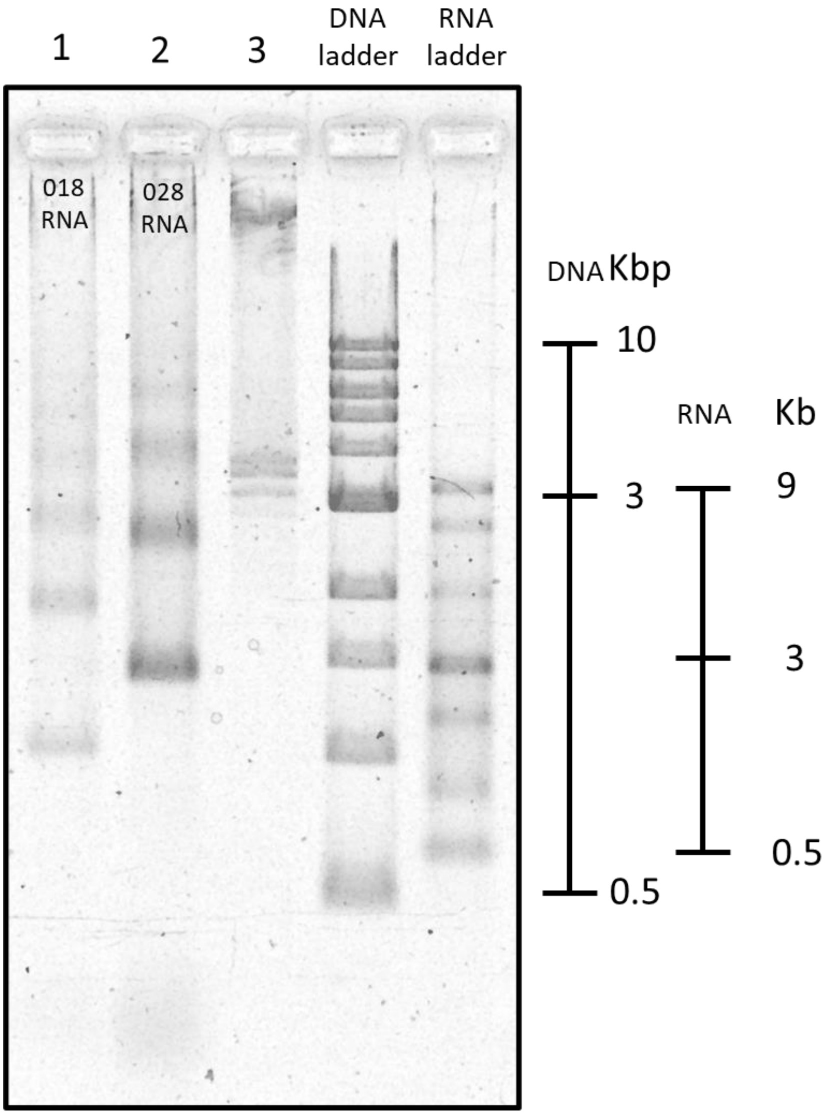


**Figure 23.** Rolling circle transcription of circular DNA constructs with different sizes and different distances between the OriC and T7RNAP promoter. The agarose gel shows a DNA ladder and ssRNA ladder on the right side. Lane -1 shows RNA product from rolling circle transcription of circular DNA construct (sequence in Supplementary Table 1) as presented in Figure 3d. Lane 2 displays RNA product from transcription of a larger DNA construct (sequence in Supplementary Table 4) with a larger separation between the OriC and promoter (~3kbp). Each band is ascribed to transcription termination at the OriC at the different transcription cycles. The Content of lane 3 is unknown. Gel: 1 % (w/v) agarose, 1 × TBE, 0.02% sodium hypochlorite.

**Figure 24.**

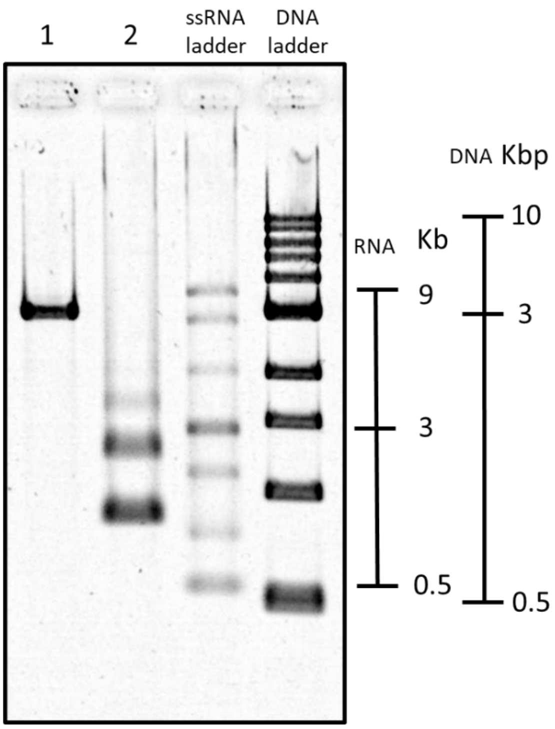


**Figure 24.** Transcription of pFGG linear DNA template. ssRNA and DNA ladder are included on the right side of the gel. Lane 1 – 2972 kbp DNA construct with no tandem repeats (sequence in Supplementary Table 5) linearized with ScaI. Lane 2 – DNase treated RNA from transcription of linear DNA in lane 1. The bottom band is ascribed to transcription until the restriction site of ScaI (1192 bp), where termination occurs as the T7RNAP falls off from the DNA template. The top band, located between 2 and 3 kb corresponds to termination at the OriC in uncut plasmids. Gel: 1 % (w/v) agarose, 1 × TBE, 0.02% sodium hypochlorite.

**Figure 25.**

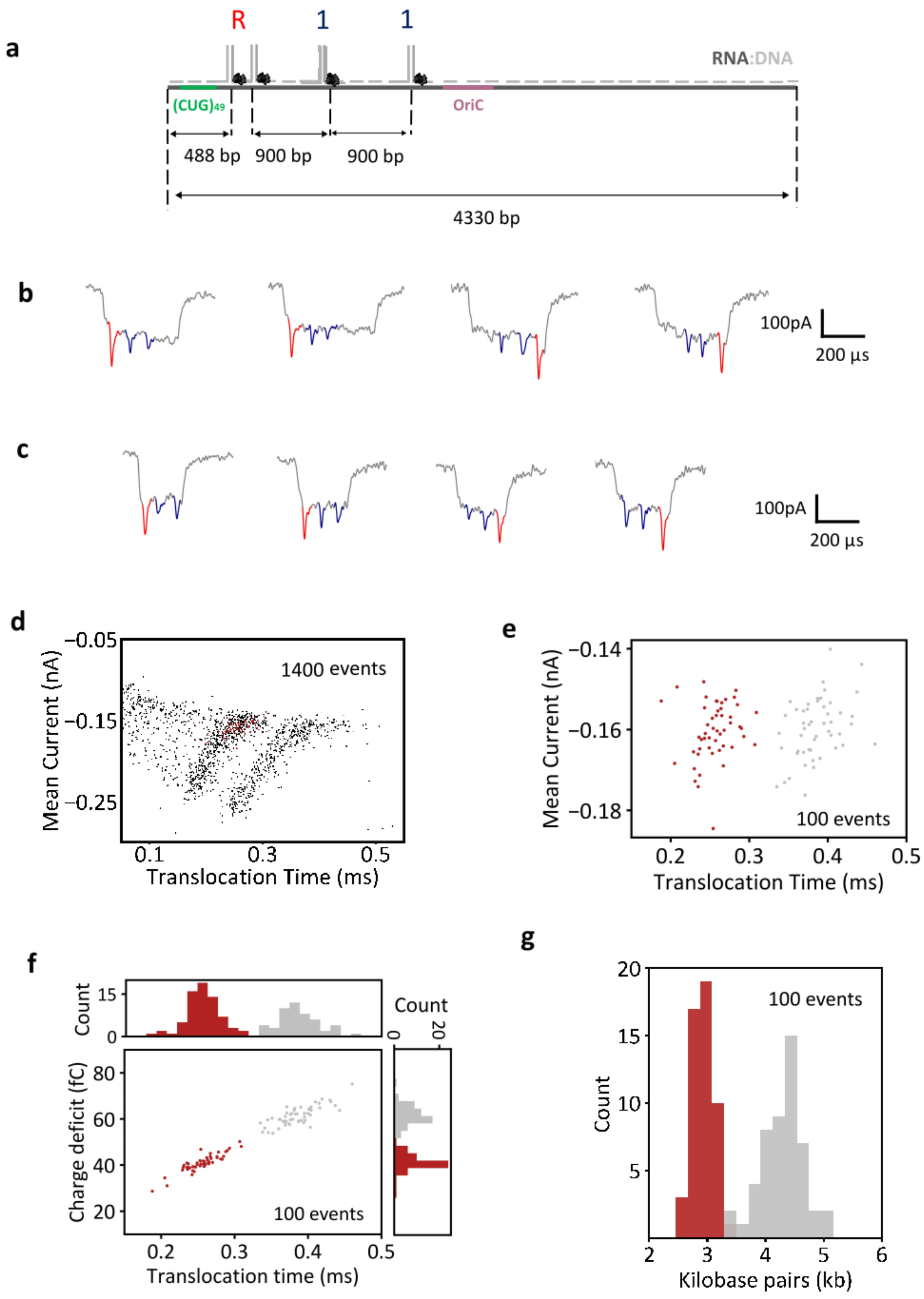
a
R
1
1
RNA:DNA
(CUG)49
OriC
488 bp
900 bp
900 bp
4330 bp
b
100pA
200 µs
c
100pA
200 µs
d
−0.05
−0.15
−0.25
Mean Current (nA)
1400 events
0.1
0.3
0.5
Translocation Time (ms)
e
−0.14
−0.16
−0.18
Mean Current (nA)
100 events
0.2
0.3
0.4
0.5
Translocation Time (ms)
f
Count
15
0
Count
0
20
Charge deficit (fC)
80
60
40
20
100 events
0.2
0.3
0.4
0.5
Translocation time (ms)
g
20
15
10
5
Count
100 events
2
3
4
5
6
Kilobase pairs (kb)

**Figure 25:** Single molecule sizing of RNA IDs produced from a 4.5 kbp DNA construct. **a** The sequence of the DNA construct is presented in Supplementary Table 4. The linearized version of the construct (DraIII) is presented Supplementary Figure 22, in lane 11, and the transcription products are shown in lane 12. The RNA ID design includes an 'R' label and two '1' bits (Supplementary Table 6). The oligos used for assembly of the hybrid are shown in supplementary Table 6. **b** Exemplary RNA ID translocation events of full-length transcripts (END). **c** Exemplary RNA ID translocation events ascribed to premature termination (PT). **d** Scatter plot of mean current against translocation time shows two distinct distributions, attributed to PT RNA IDs and END RNA IDs (from left to right). **e** Scatter plot of mean current against translocation time of 100 unfolded events. **f** Scatter plot of charge deficit against translocation time for RNA IDs of PT (red) and END (gray) RNA transcripts, which shows the linear dependence of both parameters. **g** Base pair length of molecules converted from translocation time (in f), which shows two distinct distributions. PT distribution has a mean length of (2.9 ± 0.2) kbp and END transcripts have a mean length of (4.3 ± 0.3) kbp. Errors correspond to standard deviation.

**Figure 26.**

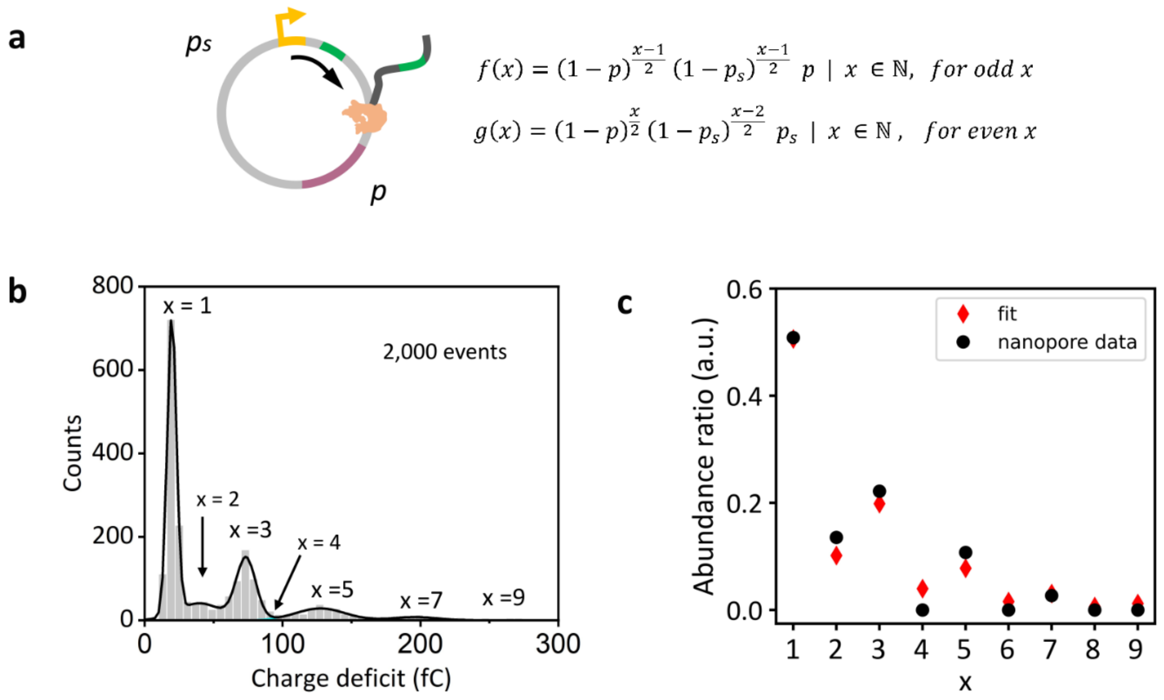


**Figure 26:** Quantitative description of T7RNAP processivity and transcription termination. **a** Considering a probability *p* of transcription terminating at the identified premature transcription termination site, and a probability $p_s$ for transcription terminating solely by dissociation of the polymerase at a different region of the plasmid. Equation *f(x)* describes the abundance of transcripts originated from premature transcription termination at OriC. Equation *g(x)* describes the transcript abundance originated from fall-off of T7RNAP at a different region of the plasmid. **b** The distribution of the charge deficit of nanopore translocation events is presented. The different transcript populations were fitted to gaussian functions, from which the relative abundance of transcripts was derived. The distributions ascribed to premature termination transcripts are labelled with *x* values of odd integers (*x* = 1, 3, 5, 7) and the minor distributions ascribed to transcripts produced from T7RNAP dissociation in a different region of the plasmid receive *x* values of even integers (*x* = 2, 4, 6, 8). **c** The relative abundance of transcripts (black) are plotted. Fitting of *f(x)* and *g(x)* is plotted in red, *p* ~ 0.51 and $p_s$ ~ 0.21, which agrees with transcription termination reported in linear DNA constructs.

**Figure 27.**

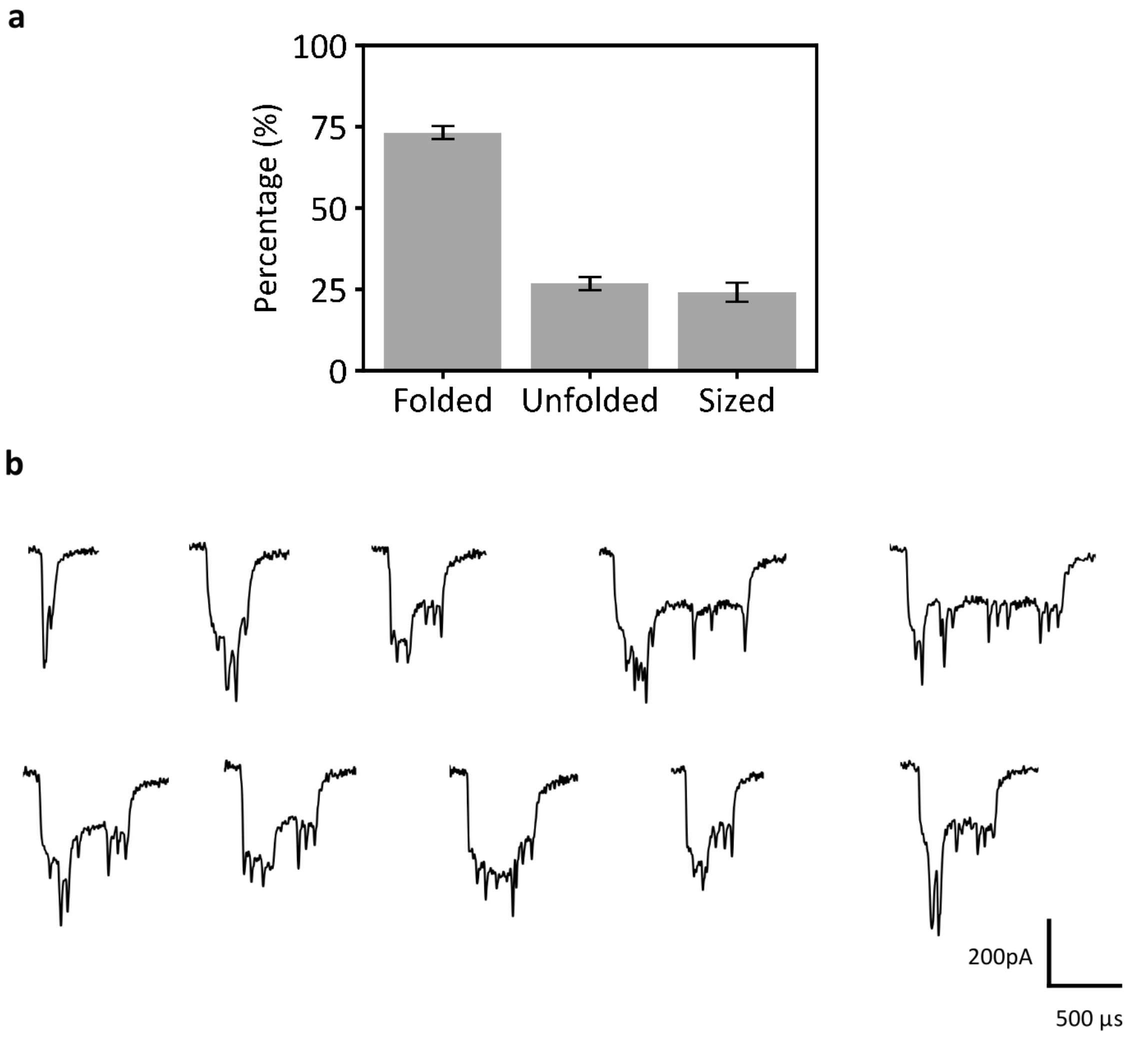


**Figure 27:** Percentage of translocation events sized. **a** Shows the percentage of folded, unfolded and sized events. The percentages were computed from 3 different nanopore measurements of RNA IDs produced from transcription of a circular DNA construct. The amount of folded events correspond to (73 ± 2)%, unfolded events constitute (27 ± 2)% of the sample, and (24 ± 3)% were sized. The errors correspond to the standard error of the mean. **b** Exemplary folded events are presented.

### Figure 28

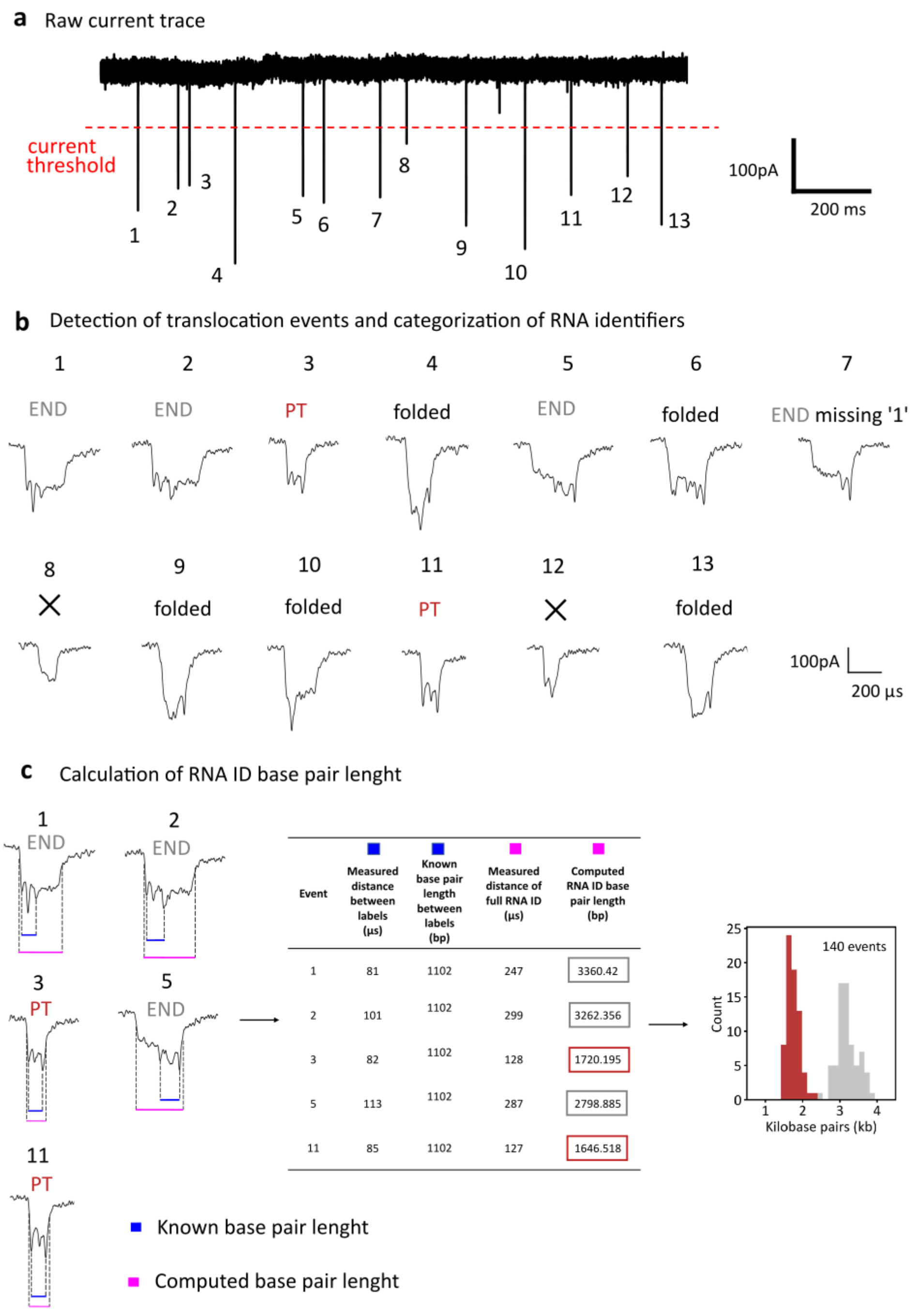


| Event | Measured distance between labels (µs) | Known base pair length between labels (bp) | Measured distance of full RNA ID (µs) | Computed RNA ID base pair length (bp) |
|---|---|---|---|---|
| 1 | 81 | 1102 | 247 | 3360.42 |
| 2 | 101 | 1102 | 299 | 3262.356 |
| 3 | 82 | 1102 | 128 | 1720.195 |
| 5 | 113 | 1102 | 287 | 2798.885 |
| 11 | 85 | 1102 | 127 | 1646.518 |

**Figure 28:** Step-by-step nanopore data analysis for RNA ID sizing. **a** An exemplary region of the ionic raw current trace is shown. We use a home-built LabVIEW code to identify translocation events by simple thresholding. Events are then analysed by using standard parameters, which include the charge deficit, minimum translocation time and mean current that are given by the length of the RNA ID molecule and its design. **b** The translocation events were then categorized into PT RNA ID, END RNA ID, folded RNA ID and translocations

associated to misfolded molecules (marked X) based on the ionic current trace of the events. **c** The base pair length of PT RNA IDs and END RNA IDs was computed using the RNA ID design. The distance between current spikes (blue) was measured (in time units). This distance corresponds to a known base pair length which was used to calculate a conversion factor. Then, the length in time units of the entire translocation event was measured (pink) and converted into a base. The base pair length was plotted as shown in the right.

**Figure 29**

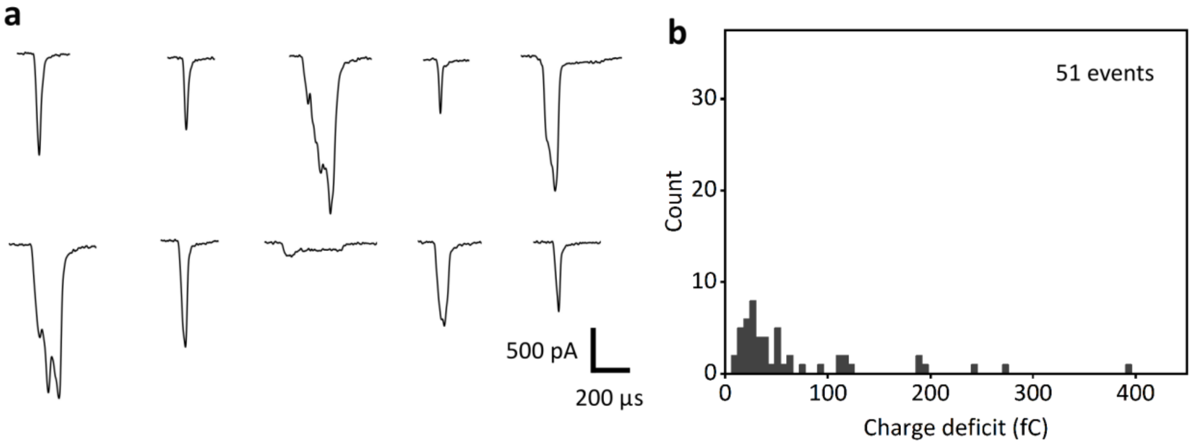


**Figure 29:** Characterization of ionic current trace shows that distinctive current traces emanate exclusively from the RNA IDs. **a** A nanopore measurement was performed in the absence of RNA IDs for 90 minutes. Translocation events detected using the following threshold parameters are shown. Minimum charge deficit: 0 fC, maximum charge deficit 400 fC, minimum translocation time: 50 µs, minimum current drop: -100 pA. Ionic current scale bar corresponds to 500 pA, and translocation time scale bar to 200 µs. **b** The charge deficit of the translocations detected are plotted, demonstrating that the current traces studied in this work originate from the translocation of RNA IDs.

#### Table 1.

Sequence of modified pJET1.2/blunt cloning vector (CloneJET PCR Cloning Kit, Thermo Fisher, Catalog number: K1231): Circular DNA (3062 bp) with inserted $(CTG)_{12}$ tandem repeats. Map of the plasmid is included at the end of the table.

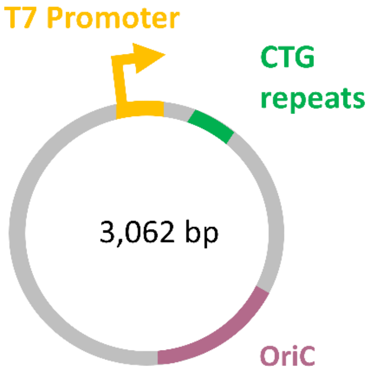


Table 1

| Sequence (5′→ 3′) |
|---|
| T7 promoter (CTG)$_{12}$ repeats OriC DraIII Cutting |
| GCCCCTGCAGCCGAATTATATTATTTTTGCCAAATAATTTTTAACAAAAGCTCTGAAGTCTTCTTC |
| ATTTAAATTCTTAGATGATACTTCATCTGGAAAATTGTCCCAATTAGTAGCAT**CACGCTGTG**AGTA |
| AGTTCTAAACCATTTTTTTATTGTTGTATTATCTCTAATCTTACTACTCGATGAGTTTTCGGTATT |
| ATCTCTATTTTTAACTTGGAGCAGGTTCCATTCATTGTTTTTTTCATCATAGTGAATAAAATCAAC |
| TGCTTTAACACTTGTGCCTGAACACCATATCCATCCGGCG**TAATACGACTCACTATAGG**GAGAGCG |
| GCCGCCAGATCTTCCGGATGGCTCGAGTTTTTCAGCAAGATCATGGTGCAGTGTAGCCGGGAATG**C** |
| **TGCTGCTGCTGCTGCTGCTGCTGCTGCTGCTGCTG**GGGGGATCACAGACCATTCTCGGCTTAAATC |
| TTTCTAGAAGATCTCCTACAATATTCTCAGCTGCCATGGAAAATCGATGTTCTTCTTTTATTCTCT |
| CAAGATTTTCAGGCTGTATATTAAAACTTATATTAAGAACTATGCTAACCACCTCATCAGGAACCG |
| TTGTAGGTGGCGTGGGTTTTCTTGGCAATCGACTCTCATGAAAACTACGAGCTAAATATTCAATAT |
| GTTCCTCTTGACCAACTTTATTCTGCATTTTTTTTGAACGAGGTTTAGAGCAAGCTTCAGGAAACT |
| GAGACAGGAATTTTATTAAAAATTTAAATTTTGAAGAAAGTTCAGGGTTAATAGCATCCATTTTTT |
| GCTTTGCAAGTTCCTCAGCATTCTTAACAAAAGACGTCTCTTTTGACATGTTTAAAGTTTAAACCT |
| CCTGTGTGAAATTATTATCCGCTCATAATTCCACACATTATACGAGCCGGAAGCATAAAGTGTAAA |
| GCCTGGGGTGCCTAATGAGTGAGCTAACTCACATTAATTGCGTTGCGCTCACTGCCAATTGCTTTC |
| CAGTCGGGAAACCTGTCGTGCCAGCTGCATTAATGAATCGGCCAACGCGCGGGGAGAGGCGGTTTG |
| CGTATTGGGCGCTCTTCCGCTTCCTCGCTCACTGACTCGCTGCGCTCGGTCGTTCGGCTGCGGCGA |
| GCGGTATCAGCTCACTCAAAGGCGGTAATACGGTTATCCACAGAATCAGGGGATAACGCAGGAAAG |
| AACATGTGAGCAAAAGGCCAGCAAAAGGCCAGGAACCGTAAAAAGGCCGCGTTGCTGGCGTT**TTTC** |
| **CATAGGCTCCGCCCCCCTGACGAGCATCACAAAAATCGACGCTCAAGTCAGAGGTGGCGAAACCCG** |
| **ACAGGACTATAAAGATACCAGGCGTTTCCCCCTGGAAGCTCCCTCGTGCGCTCTCCTGTTCCGACC** |
| **CTGCCGCTTACCGGATACCTGTCCGCCTTTCTCCCTTCGGGAAGCGTGGCGCTTTCTCATAGCTCA** |

**CGCTGTAGGTATCTCAGTTCGGTGTAGGTCGTTCGCTCCAAGCTGGGCTGTGTGCACGAACCCCCC**
**GTTCAGCCCGACCGCTGCGCCTTATCCGGTAACTATCGTCTTGAGTCCAACCCGGTAAGACACGAC**
**TTATCGCCACTGGCAGCAGCCACTGGTAACAGGATTAGCAGAGCGAGGTATGTAGGCGGTGCTACA**
**GAGTTCTTGAAGTGGTGGCCTAACTACGGCTACACTAGAAGGACAGTATTTGGTATCTGCGCTCTG**
**CTGAAGCCAGTTACCTTCGGAAAAAGAGTTGGTAGCTCTTGATCCGGCAAACAAACCACCGCTGGT**
**AGCGGTGGTTTTTTTGTTTGCAAGCAGCAGATTACGCGCAGAAAAAAAGGATCTCAA**GAAGATCCT
TTGATCTTTTCTACGGGGTCTGACGCTCAGTGGAACGAAAACTCACGTTAAGGGATTTTGGTCATG
AGATTATCAAAAAGGATCTTCACCTAGATCCTTTTAAATTAAAAATGAAGTTTTAAATCAATCTAA
AGTATATATGAGTAAACTTGGTCTGACAGTTACCAATGCTTAATCAGTGAGGCACCTATCTCAGCG
ATCTGTCTATTTCGTTCATCCATAGTTGCCTGACTCCCCGTCGTGTAGATAACTACGATACGGGAG
GGCTTACCATCTGGCCCCAGTGCTGCAATGATACCGCGAGACCCACGCTCACCGGCTCCAGATTTA
TCAGCAATAAACCAGCCAGCCGGAAGGGCCGAGCGCAGAAGTGGTCCTGCAACTTTATCCGCCTCC
ATCCAGTCTATTAATTGTTGCCGGGAAGCTAGAGTAAGTAGTTCGCCAGTTAATAGTTTGCGCAAC
GTTGTTGCCATTGCTACAGGCATCGTGGTGTCACGCTCGTCGTTTGGTATGGCTTCATTCAGCTCC
GGTTCCCAACGATCAAGGCGAGTTACATGATCCCCCATGTTGTGCAAAAAAGCGGTTAGCTCCTTC
GGTCCTCCGATCGTTGTCAGAAGTAAGTTGGCCGCAGTGTTATCACTCATGGTTATGGCAGCACTG
CATAATTCTCTTACTGTCATGCCATCCGTAAGATGCTTTTCTGTGACTGGTGAGTACTCAACCAAG
TCATTCTGAGAATAGTGTATGCGGCGACCGAGTTGCTCTTGCCCGGCGTCAATACGGGATAATACC
GCGCCACATAGCAGAACTTTAAAAGTGCTCATCATTGGAAAACGTTCTTCGGGGCGAAAACTCTCA
AGGATCTTACCGCTGTTGAGATCCAGTTCGATGTAACCCACTCGTGCACCCAACTGATCTTCAGCA
TCTTTTACTTTCACCAGCGTTTCTGGGTGAGCAAAAACAGGAAGGCAAAATGCCGCAAAAAAGGGA
ATAAGGGCGACACGGAAATGTTGAATACTCATACTCTTCCTTTTTCAATATTATTGAAGCATTTAT
CAGGGTTATTGTCTCATGAGCGGATACATATTTGAATGTATTTAGAAAAATAAACAAATAGGGGTT
CCGCGCACATTTCCCCGAAAAGTGCCACCTGACGTCTAAGAAACCATTATTATCATGACATTAACC
TATAAAAATAGGCGTATCACGAGGCC

Map of the plasmid showing restriction sites, T7RNAP promoter sequence, the OriC and the CTG repeats.

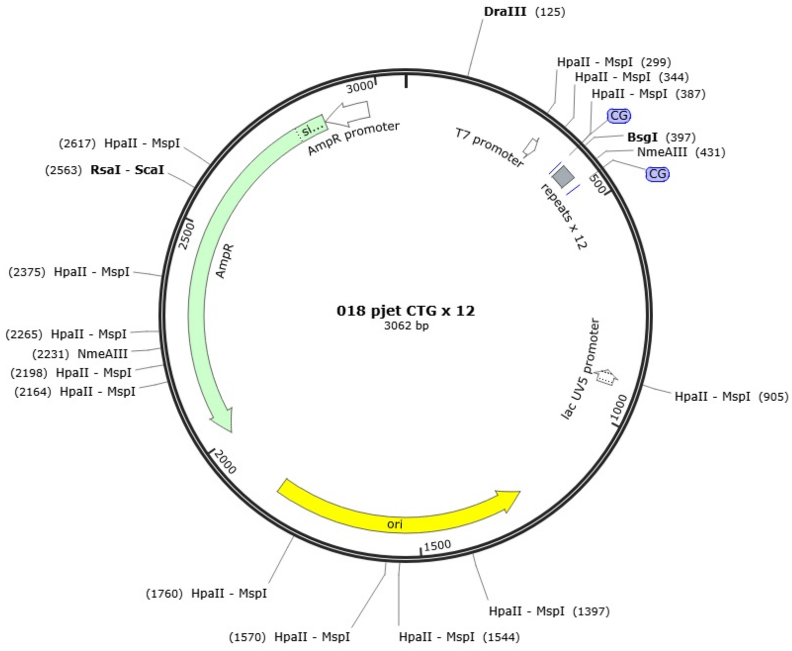

#### Table 2.

Sequence of oligonucleotides in RNA ID design for characterization of transcripts from linear DNA. This design is used for the study of RNA from premature termination (PT) and RNA from full transcription of the linear template (Figure 1). The oligos are complementary to the region of the circular plasmid (Supplementary Table 1) covering from the promoter to DraIII restriction site.

▲ Complementary DNA oligo ▲ Bit '1' oligo ▲ Repeats label

Table 2

| Oligo number | Sequence (5'→ 3') |
|---|---|
| 1 | AGCGTGATGCTACTAATTGGGACAATTTTCCAGATGAAGT |
| 2 | ATCATCTAAGAATTTAAATGAAGAAGACTTCAGAGCTTTT |
| 3 | GTTAAAAATTATTTGGCAAAAATAATATAATTCGGCTGCA |
| 4 | GGGGCGGCCTCGTGATACGCCTATTTTTATAGGTTAATGT |
| 5 | CATGATAATAATGGTTTCTTAGACGTCAGGTGGCACTTTT |
| 6 | CGGGGAAATGTGCGCGGAACCCCTATTTGTTTATTTTTCT |
| 7 | AAATACATTCAAATATGTATCCGCTCATGAGACAATAACC |
| 8 | CTGATAAATGCTTCAATAATATTGAAAAAGGAAGAGTATG |
| 9 | AGTATTCAACATTTCCGTGTCGCCCTTATTCCCTTTTTTG |
| 10 | CGGCATTTTGCCTTCCTGTTTTTGCTCACCCAGAAACGCT |
| 11 | GGTGAAAGTAAAAGATGCTGAAGATC |
| 12 | AGTTGGGTGCACGAGTGGGTTACATCG |
| 13 | AACTGGATCTCAACAGCGGTAAGAT |
| 14 | CCTTGAGAGTTTTCGCCCCGAAGAACGTTTTCCAATGATG |
| 15 | AGCACTTTTAAAGTTCTGCTATGTGGCGCGGTATTATCCC |
| 16 | GTATTGACGCCGGGCAAGAGCAACTCGGTCGCCGCATACA |
| 17 | CTATTCTCAGAATGACTTGGTTGAGTACTCACCAGTCACA |
| 18 | GAAAAGCATCTTACGGATGGCATGACAGTAAGAGAATTAT |
| 19 | GCAGTGCTGCCATAACCATGAGTGATAACACTGCGGCCAA |
| 20 | CTTACTTCTGACAACGATCGGAGGACCGAAGGAGCTAACC |
| 21 | GCTTTTTTGCACAACATGGGGGATCATGTAACTCGCCTTG |
| 22 | ATCGTTGGGAACCGGAGCTGAATGAAGCCATACCAAACGA |
| 23 | CGAGCGTGACACCACGATGCCTGTAGCAATGGCAACAACG |
| 24 | TTGCGCAAACTATTAACTGGCGAACTACTTACTCTAGCTT |
| 25 | CCCGGCAACAATTAATAGACTGGATGGAGGCGGATAAAGT |
| 26 | TGCAGGACCACTTCTGCGCTCGGCCCTTCCGGCTGGCTGG |
| 27 | TTTATTGCTGATAAATCTGGAGCCGGT |
| 28 | GAGCGTGGGTCTCGCGGTATCATTGCAG |

| | |
|---|---|
| 29 | CACTGGGGCCAGATGGTAAGCCCTC |
| 30 | CCGTATCGTAGTTATCTACACGACGGGGAGTCAGGCAACT |
| 31 | ATGGATGAACGAAATAGACAGATCGCTGAGATAGGTGCCT |
| 32 | CACTGATTAAGCATTGGTAACTGTCAGACCAAGTTTACTC |
| 33 | ATATATACTTTAGATTGATTTAAAACTTCATTTTTAATTT |
| 34 | AAAAGGATCTAGGTGAAGATCCTTTTTGATAATCTCATGA |
| 35 | CCAAAATCCCTTAACGTGAGTTTTCGTTCCACTGAGCGTC |
| 36 | AGACCCCGTAGAAAAGATCAAAGGATCTTCTTGAGATCCT |
| 37 | TTTTTTCTGCGCGTAATCTGCTGCTTGCAAACAAAAAAAC |
| 38 | CACCGCTACCAGCGGTGGTTTGTTTGCCGGATCAAGAGCT |
| 39 | ACCAACTCTTTTTCCGAAGGTAACTGGCTTCAGCAGAGCG |
| 40 | CAGATACCAAATACTGTCCTTCTAGTGTAGCCGTAGTTAG |
| 41 | GCCACCACTTCAAGAACTCTGTAGCACCGCCTACATACCT |
| 42 | CGCTCTGCTAATCCTGTTACCAGTGGCTGCTGCCAGTGGC |
| 43 | GATAAGTCGTGTCTTACCGGGTTGGAC |
| 44 | TCAAGACGATAGTTACCGGATAAGGCGC |
| 45 | AGCGGTCGGGCTGAACGGGGGGTTCTTTGGATATCACTCATTAGTGGT |
| 46 | GTGCACACAGCCCAGCTTGGAGCGAACGACCTACACCGAA |
| 47 | CTGAGATACCTACAGCGTGAGCTATGAGAAAGCGCCACGC |
| 48 | TTCCCGAAGGGAGAAAGGCGGACAGGTATCCGGTAAGCGG |
| 49 | CAGGGTCGGAACAGGAGAGCGCACGAGGGAGCTTCCAGGG |
| 50 | GGAAACGCCTGGTATCTTTATAGTCCTGTCGGGTTTCGCC |
| 51 | ACCTCTGACTTGAGCGTCGATTTTTGTGATGCTCGTCAGG |
| 52 | GGGGCGGAGCCTATGGAAAAACGCCAGCAACGCGGCCTTT |
| 53 | TTACGGTTCCTGGCCTTTTGCTGGCCTTTTGCTCACATGT |
| 54 | TCTTTCCTGCGTTATCCCCTGATTCTGTGGATAACCGTAT |
| 55 | TACCGCCTTTGAGTGAGCTGATACCGCTCGCCGCAGCCGA |
| 56 | ACGACCGAGCGCAGCGAGTCAGTGAGCGAGGAAGCGGAAG |
| 57 | AGCGCCCAATACGCAAACCGCCTCTCCCCGCGCGTTGGCC |
| 58 | GATTCATTAATGCAGCTGGCACGACAGGTTTCCCGACTGG |
| 59 | AAAGCAATTGGCAGTGAGCGCAACGCA |
| 60 | ATTAATGTGAGTTAGCTCACTCATTAGG |
| 61 | CACCCCAGGCTTTACACTTTATGCTTTTGGATATCACTCATTAGTGGT |
| 62 | TCCGGCTCGTATAATGTGTGGAATTATGAGCGGATAATAA |
| 63 | TTTCACACAGGAGGTTTAAACTTTAAACATGTCAAAAGAG |
| 64 | ACGTCTTTTGTTAAGAATGCTGAGGAACTTGCAAAGCAAA |
| 65 | AAATGGATGCTATTAACCCTGAACTTTCTTCAAAATTTAA |
| 66 | ATTTTTAATAAAATTCCTGTCTCAGTTTCCTGAAGCTTGC |
| 67 | TCTAAACCTCGTTCAAAAAAAAATGCAGAATAAAGTTGGTC |
| 68 | AAGAGGAACATATTGAATATTTAGCTCGTAGTTTTCATGA |
| 69 | GAGTCGATTGCCAAGAAAACCCACGCCACCTACAACGGTT |
| 70 | CCTGATGAGGTGGTTAGCATAGTTCTTAATATAAGTTTTA |
| 71 | ATATACAGCCTGAAAATCTTGAGAGAATAAAAGAAGAACA |
| 72 | TCGATTTTCCATGGCAGCTGAGAATATTGTAGGAGATCTT |
| 73 | CTAGAAAGATTTAAGCCGAGAATGGTCTGTGATCCCCC |
| 74 | CATTCCCGGCTACACTGCACCATGATCTTGCTGAAAAACT |

| | |
|---|---|
| **75** | CGAGCCATCCGGAAGATCTGGCGGCCGCTCTCCC |
| **76** | CAGCAGCAGCAGCAGCAGCAGCAGCAGCAGTTTGGATATCACTCATTAGTGGT/3’-biotin/ |

#### Table 3.

Sequence of oligonucleotides in RNA ID design for characterization of transcripts from circular DNA. This design is used for the study of RNA IDs from multiple transcription cycles (Figure 3). The oligos are complementary to the entire circular plasmid (Supplementary Table 1). This table includes the same oligonucleotides as Table 2, however it contains 5 extra oligonucelotides (76, 77, 78, 79, 80) which are complementary to the region extending from DraIII restriction site to the T7RNAP promoter.

▲ Complementary DNA oligo ▲ Bit '1' oligo ▲ Repeats label

Table 3

| Oligo number | Sequence (5′→ 3′) |
|---|---|
| 1 | AGCGTGATGCTACTAATTGGGACAATTTTCCAGATGAAGT |
| 2 | ATCATCTAAGAATTTAAATGAAGAAGACTTCAGAGCTTTT |
| 3 | GTTAAAAATTATTTGGCAAAAATAATATAATTCGGCTGCA |
| 4 | GGGGCGGCCTCGTGATACGCCTATTTTTATAGGTTAATGT |
| 5 | CATGATAATAATGGTTTCTTAGACGTCAGGTGGCACTTTT |
| 6 | CGGGGAAATGTGCGCGGAACCCCTATTTGTTTATTTTTCT |
| 7 | AAATACATTCAAATATGTATCCGCTCATGAGACAATAACC |
| 8 | CTGATAAATGCTTCAATAATATTGAAAAAGGAAGAGTATG |
| 9 | AGTATTCAACATTTCCGTGTCGCCCTTATTCCCTTTTTTG |
| 10 | CGGCATTTTGCCTTCCTGTTTTTGCTCACCCAGAAACGCT |
| 11 | GGTGAAAGTAAAAGATGCTGAAGATC |
| 12 | AGTTGGGTGCACGAGTGGGTTACATCG |
| 13 | AACTGGATCTCAACAGCGGTAAGAT |
| 14 | CCTTGAGAGTTTTCGCCCCGAAGAACGTTTTCCAATGATG |
| 15 | AGCACTTTTAAAGTTCTGCTATGTGGCGCGGTATTATCCC |
| 16 | GTATTGACGCCGGGCAAGAGCAACTCGGTCGCCGCATACA |
| 17 | CTATTCTCAGAATGACTTGGTTGAGTACTCACCAGTCACA |
| 18 | GAAAAGCATCTTACGGATGGCATGACAGTAAGAGAATTAT |
| 19 | GCAGTGCTGCCATAACCATGAGTGATAACACTGCGGCCAA |
| 20 | CTTACTTCTGACAACGATCGGAGGACCGAAGGAGCTAACC |
| 21 | GCTTTTTTGCACAACATGGGGGATCATGTAACTCGCCTTG |
| 22 | ATCGTTGGGAACCGGAGCTGAATGAAGCCATACCAAACGA |
| 23 | CGAGCGTGACACCACGATGCCTGTAGCAATGGCAACAACG |
| 24 | TTGCGCAAACTATTAACTGGCGAACTACTTACTCTAGCTT |
| 25 | CCCGGCAACAATTAATAGACTGGATGGAGGCGGATAAAGT |
| 26 | TGCAGGACCACTTCTGCGCTCGGCCCTTCCGGCTGGCTGG |
| 27 | TTTATTGCTGATAAATCTGGAGCCGGT |

| | |
|---|---|
| **28** | GAGCGTGGGTCTCGCGGTATCATTGCAG |
| **29** | CACTGGGGCCAGATGGTAAGCCCTC |
| **30** | CCGTATCGTAGTTATCTACACGACGGGGAGTCAGGCAACT |
| **31** | ATGGATGAACGAAATAGACAGATCGCTGAGATAGGTGCCT |
| **32** | CACTGATTAAGCATTGGTAACTGTCAGACCAAGTTTACTC |
| **33** | ATATATACTTTAGATTGATTTAAAACTTCATTTTTAATTT |
| **34** | AAAAGGATCTAGGTGAAGATCCTTTTTGATAATCTCATGA |
| **35** | CCAAAATCCCTTAACGTGAGTTTTCGTTCCACTGAGCGTC |
| **36** | AGACCCCGTAGAAAAGATCAAAGGATCTTCTTGAGATCCT |
| **37** | TTTTTTCTGCGCGTAATCTGCTGCTTGCAAACAAAAAAAC |
| **38** | CACCGCTACCAGCGGTGGTTTGTTTGCCGGATCAAGAGCT |
| **39** | ACCAACTCTTTTTCCGAAGGTAACTGGCTTCAGCAGAGCG |
| **40** | CAGATACCAAATACTGTCCTTCTAGTGTAGCCGTAGTTAG |
| **41** | GCCACCACTTCAAGAACTCTGTAGCACCGCCTACATACCT |
| **42** | CGCTCTGCTAATCCTGTTACCAGTGGCTGCTGCCAGTGGC |
| **43** | GATAAGTCGTGTCTTACCGGGTTGGAC |
| **44** | TCAAGACGATAGTTACCGGATAAGGCGC |
| **45** | AGCGGTCGGGCTGAACGGGGGGTTCTTTGGATATCACTCATTAGTGGT |
| **46** | GTGCACACAGCCCAGCTTGGAGCGAACGACCTACACCGAA |
| **47** | CTGAGATACCTACAGCGTGAGCTATGAGAAAGCGCCACGC |
| **48** | TTCCCGAAGGGAGAAAGGCGGACAGGTATCCGGTAAGCGG |
| **49** | CAGGGTCGGAACAGGAGAGCGCACGAGGGAGCTTCCAGGG |
| **50** | GGAAACGCCTGGTATCTTTATAGTCCTGTCGGGTTTCGCC |
| **51** | ACCTCTGACTTGAGCGTCGATTTTTGTGATGCTCGTCAGG |
| **52** | GGGGCGGAGCCTATGGAAAAACGCCAGCAACGCGGCCTTT |
| **53** | TTACGGTTCCTGGCCTTTTGCTGGCCTTTTGCTCACATGT |
| **54** | TCTTTCCTGCGTTATCCCCTGATTCTGTGGATAACCGTAT |
| **55** | TACCGCCTTTGAGTGAGCTGATACCGCTCGCCGCAGCCGA |
| **56** | ACGACCGAGCGCAGCGAGTCAGTGAGCGAGGAAGCGGAAG |
| **57** | AGCGCCCAATACGCAAACCGCCTCTCCCCGCGCGTTGGCC |
| **58** | GATTCATTAATGCAGCTGGCACGACAGGTTTCCCGACTGG |
| **59** | AAAGCAATTGGCAGTGAGCGCAACGCA |
| **60** | ATTAATGTGAGTTAGCTCACTCATTAGG |
| **61** | CACCCCAGGCTTTACACTTTATGCTTTTGGATATCACTCATTAGTGGT |
| **62** | TCCGGCTCGTATAATGTGTGGAATTATGAGCGGATAATAA |
| **63** | TTTCACACAGGAGGTTTAAACTTTAAACATGTCAAAAGAG |
| **64** | ACGTCTTTTGTTAAGAATGCTGAGGAACTTGCAAAGCAAA |
| **65** | AAATGGATGCTATTAACCCTGAACTTTCTTCAAAATTTAA |
| **66** | ATTTTTAATAAAATTCCTGTCTCAGTTTCCTGAAGCTTGC |
| **67** | TCTAAACCTCGTTCAAAAAAAATGCAGAATAAAGTTGGTC |
| **68** | AAGAGGAACATATTGAATATTTAGCTCGTAGTTTTCATGA |
| **69** | GAGTCGATTGCCAAGAAAACCCACGCCACCTACAACGGTT |
| **70** | CCTGATGAGGTGGTTAGCATAGTTCTTAATATAAGTTTTA |
| **71** | ATATACAGCCTGAAAATCTTGAGAGAATAAAAGAAGAACA |
| **72** | TCGATTTTCCATGGCAGCTGAGAATATTGTAGGAGATCTT |
| **73** | CTAGAAAGATTTAAGCCGAGAATGGTCTGTGATCCCCC |

| | |
|---|---|
| **74** | CATTCCCGGCTACACTGCACCATGATCTTGCTGAAAAACT |
| **75** | CGAGCCATCCGGAAGATCTGGCGGCCGCTCTCCC |
| **76** | TATAGTGAGTCGTATTACGCCGGATGGATATGGTGTTC |
| **77** | AGGCACAAGTGTTAAAGCAGTTGATTTTATTCACTATG |
| **78** | ATGAAAAAAACAATGAATGGAACCTGCTCCAAGTTA |
| **79** | AAAATAGAGATAATACCGAAAACTCATCGAGTAGTA |
| **80** | AGATTAGAGATAATACAACAATAAAAAAATGGTTTAGAACTTACTCACAGC |
| **81** | CAGCAGCAGCAGCAGCAGCAGCAGCAGCAGTTTGGATATCACTCATTAGTGGT/3’-biotin/ |

#### Table 4.

Sequence of modified pJET1.2/blunt cloning vector (CloneJET PCR Cloning Kit, Thermo Fisher, Catalog number: K1231): Circular DNA (4527 bp) with inserted $(CTG)_{49}$ tandem repeats.  The sequence of the plasmid was verified by whole plasmid sequencing. Map of the plasmid is included at the end of the table.

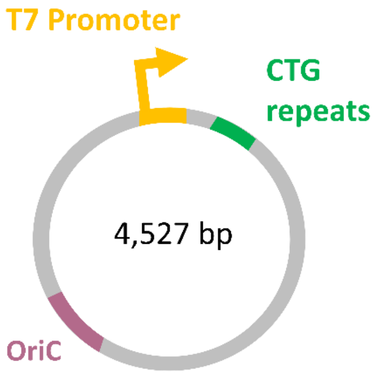


Table 4

| Sequence (5′→ 3′) |
| --- |
| T7 promoter $(CTG)_{49}$ repeats OriC DraIII Cutting |
| GCATCCATTTTTTGCTTTGCAAGTTCCTCAGCATTCTTAACAAAAGACGTCTCTTTTGACATGTTT<br>AAAGTTTAAACCTCCTGTGTGAAATTATTATCCGCTCATAATTCCACACATTATACTGAGAGATCC<br>CCTCATAATTTCCCCAAAGCGTAACCATGTGTGAATAAATTTTGAGCTAGTAGGGTTGCAGCCACG<br>AGTAAGTCTTCCCTTGTTATTGTGTAGCCAGAATGCCGCAAAACTTCCATGCCTAAGCGAACTGTT<br>GAGAGTACGTTTCGATTTCTGACTGTGTTAGCCTGGAAGTGCTTGTCCCAACCTTGTTTCTGAGCA<br>TGAACGCCCGCAAGCCAACATGTTAGTTGAAGCATCAGGGCGATTAGCAGCATGATATCAAAACGC<br>TCTGAGCTGCTCGTTCGGCTATGGCGTAGGCCTAGTCCGTAGGCAGGACTTTTCAAGTCTCGGAAG<br>GTTTCTTCAATCTGCATTCGCTTCGAATAGATATTAACAAGTTGTTTGGGTGTTCGAATTTCAACA<br>GGTAAGTTAGTTGCTAGAACCCATGGCTCCTTTGCCGACGCTGAGTAGATTTTAGGTGACGGGTGG<br>TGACAATGAGTCCGTGTCGAGCGCTGATTTTTTCGGCCTTTAGAGCGAGATTTATACAATAGAATT<br>TGGCATGAGATTGGATTGCTTTTAGTCAGCCTCTTATAGCCTAAAGTCTTTGAGTGACTAGATGAC<br>ATATCATGTAAGTTGCTGATAGGTTTCCAGTTTTTCCGCTCCTAGGTCTGCATATTGTACTTTTCC<br>TCTTACTCGACTTAACCAGTACCAACCCAGCTTCTCAACGGATTTATACCATGGCACTTTAAAGCC<br>AGCATCACTGACAATGAGCGGTGTGGTGTTACTCGGTAGAATGCTCGCAAGGTCGGCTAGAAATTG<br>GTCATGAGCTTTCTTTGAACATTGCTCTGAAAGCGGGAACGCTTTCTCATAAAGAGTAACAGAACG<br>ACCGTGTAGTGCGACTGAAGCTCGCAATACCATAAGTCGTTTTTGCTCACGAATATCAGACCAGTC<br>AACAAGTACAATGGGCATCGTATTGCCCGAACAGATAAAGCTAGCATGCCAACGGTATACAGCGAG<br>TCGCTCTTTGTGGAGGTGACGATTACCTAACAATCGGTCGATTCGTTTGATGTTATGTTTTGTTCT<br>CGCTTTGGTTGGCAGGTTACGGCCAAGTTCGGTAAGAGTGAGAGTTTTACAGTCAAGTAATGCGTG<br>GCAAGCCAACGTTAAGCTGTTGAGTCGTTTTAAGTGTAATTCGGGGCAGAATTGGTAAAGAGAGTC<br>GTGTAAAATATCGAGTTCGCACATCTTGTTGTCTGATTATTGATTTTTCGCGAAACCATTTGATCA |

TATGACAAGATGTGTATCCACCTTAACTTAATGATTTTTACCAAAATCATTAGGGGATTCATCAGC
ACATTATACGAGCCGGAAGCATAAAGTGTAAAGCCTGGGGTGCCTAATGAGTGAGCTAACTCACAT
TAATTGCGTTGCGCTCACTGCCAATTGCTTTCCAGTCGGGAAACCTGTCGTGCCAGCTGCATTAAT
GAATCGGCCAACGCGCGGGGAGAGGCGGTTTGCGTATTGGGCGCTCTTCCGCTTCCTCGCTCACTG
ACTCGCTGCGCTCGGTCGTTCGGCTGCGGCGAGCGGTATCAGCTCACTCAAAGGCGGTAATACGGT
TATCCACAGAATCAGGGGATAACGCAGGAAAGAACATGTGAGCAAAAGGCCAGCAAAAGGCCAGGA
ACCGTAAAAAGGCCGCGTTGCTGGCGTT**TTTCCATAGGCTCCGCCCCCCTGACGAGCATCACAAAA**
**ATCGACGCTCAAGTCAGAGGTGGCGAAACCCGACAGGACTATAAAGATACCAGGCGTTTCCCCCTG**
**GAAGCTCCCTCGTGCGCTCTCCTGTTCCGACCCTGCCGCTTACCGGATACCTGTCCGCCTTTCTCC**
**CTTCGGGAAGCGTGGCGCTTTCTCATAGCTCACGCTGTAGGTATCTCAGTTCGGTGTAGGTCGTTC**
**GCTCCAAGCTGGGCTGTGTGCACGAACCCCCCGTTCAGCCCGACCGCTGCGCCTTATCCGGTAACT**
**ATCGTCTTGAGTCCAACCCGGTAAGACACGACTTATCGCCACTGGCAGCAGCCACTGGTAACAGGA**
**TTAGCAGAGCGAGGTATGTAGGCGGTGCTACAGAGTTCTTGAAGTGGTGGCCTAACTACGGCTACA**
**CTAGAAGAACAGTATTTGGTATCTGCGCTCTGCTGAAGCCAGTTACCTTCGGAAAAAGAGTTGGTA**
**GCTCTTGATCCGGCAAACAAACCACCGCTGGTAGCGGTGGTTTTTTTGTTTGCAAGCAGCAGATTA**
**CGCGCAGAAAAAAAGGATCTCAA**GAAGATCCTTTGATCTTTTCTACGGGGTCTGACGCTCAGTGGA
ACGAAAACTCACGTTAAGGGATTTTGGTCATGAGATTATCAAAAAGGATCTTCACCTAGATCCTTT
TAAATTAAAAATGAAGTTTTAAATCAATCTAAAGTATATATGAGTAAACTTGGTCTGACAGTTACC
AATGCTTAATCAGTGAGGCACCTATCTCAGCGATCTGTCTATTTCGTTCATCCATAGTTGCCTGAC
TCCCCGTCGTGTAGATAACTACGATACGGGAGGGCTTACCATCTGGCCCCAGTGCTGCAATGATAC
CGCGAGACCCACGCTCACCGGCTCCAGATTTATCAGCAATAAACCAGCCAGCCGGAAGGGCCGAGC
GCAGAAGTGGTCCTGCAACTTTATCCGCCTCCATCCAGTCTATTAATTGTTGCCGGGAAGCTAGAG
TAAGTAGTTCGCCAGTTAATAGTTTGCGCAACGTTGTTGCCATTGCTACAGGCATCGTGGTGTCAC
GCTCGTCGTTTGGTATGGCTTCATTCAGCTCCGGTTCCCAACGATCAAGGCGAGTTACATGATCCC
CCATGTTGTGCAAAAAAGCGGTTAGCTCCTTCGGTCCTCCGATCGTTGTCAGAAGTAAGTTGGCCG
CAGTGTTATCACTCATGGTTATGGCAGCACTGCATAATTCTCTACTGTCATGCCATCCGTAAGATG
CTTTTCTGTGACTGGTGAGTACTCAACCAAGTCATTCTGAGAATAGTGTATGCGGCGACCGAGTTG
CTCTTGCCCGGCGTCAATACGGGATATAACCGCGCCACAGTAGCAGAGCTTTAAAAGTGCTCATCA
TTGGAAAACGTTCTTCGGGGCGGAAAACTCTTCAAGGATCTTACCGCTGTTGAGATCCAGTTCGAT
GTAACCCACTCGTGAACCGAACTGATCTTCGTGCATATTTTACCGCCTTTCCCAGCGTTTCTGGGT
CAAACCTTCTTAAAAACAGGAAGGCAAAATGCCGCAAAAAAGGGAATAAGGGCGACACGGAAATGT
TGAATACTCATACTCTTCCTTTTTCAATATTATTGAAGCATTTATCAGGGTTATTGTCTCATGAGC
GGATACATATTTGAATGTATTTAGAAAAATAAACAAATAGGGGTTCCGCGCACATTTCCCCGAAAA
GTGCCACCTGACGTCTAAGAAACCATTATTATCATGACATTAACCTATAAAAATAGGCGTATCACG
AGGCCGCCCCTGCAGCCGAATTATATTATTTTTGCCAAATAATTTTTAACAAAAGCTCTGAAGTCT
TCTTCATTTAAATTCTTAGATGATACTTCATCTGGAAAATTGTCCCAATTAGTAGCAT**CACGCTGT**
**G**AGTAAGTTCTAAACCATTTTTTTTATTGTTGTATTATCTCTAATCTTACTACTCGATGAGTTTTC
GGTATTATCTCTATTTTTAACTTGGAGCAGGTTCCATTCATTGTTTTTTTCATCATAGTGAATAAA
ATCAACTGCTTTAACACTTGTGCCTGAACACCATATCCATCCGGCG**TAATACGACTCACTATAGG**G
AGAGCGGCCGCCAGATCTTCCGGATGGCTCGAGTTTTCAGCAAGATCATGGTGCAGTGTAGCCGG
GAATG**CTGCTGCTGCTGCTGCTGCTGCTGCTGCTGCTGCTGCTGCTGCTGCTGCTGCTGCTGC**
**TGCTGCTGCTGCTGCTGCTGCTGCTGCTGCTGCTGCTGCTGCTGCTGCTGCTGCTGCTGCTGC**
**TGCTGCTGCTGCTGCTG**CTGGGGGGGATCACAGACCATTCTCGGCTTAAATCTTTCTAGAAGATCT
CCTACAATATTCTCAGCTGCCATGGAAAATCGATGTTCTTCTTTTATTCTCTCAAGATTTTCAGGC
TGTATATTAAAACTTATATTAAGAACTATGCTAACCACCTCATCAGGAACCGTTGTAGGTGGCGTG
GGTTTTCTTGGCAATCGACTCTCATGAAAACTACGAGCTAAATATTCAATATGTTCCTCTTGACCA
ACTTTATTCTGCATTTTTTTTGAACGAGGTTTAGAGCAAGCTTCAGGAAACTGAGACAGGAATTTT
ATTAAAAATTTAAATTTTGAAGAAAGTTCAGGGTTAATA

Map of the plasmid showing restriction sites, T7RNAP promoter sequence, the OriC and the CTG repeats.

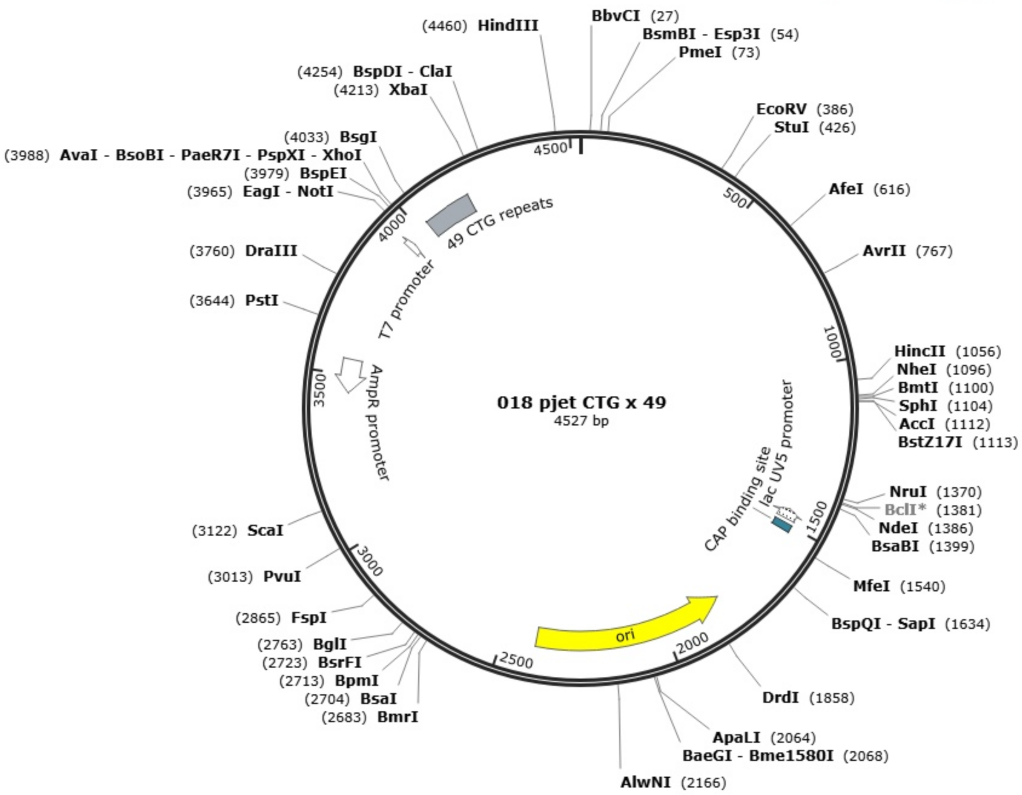

**Table 5.**

PFGG plasmid. The plasmid was produced by IDT and verified using next-generation sequencing and Sanger sequencing.

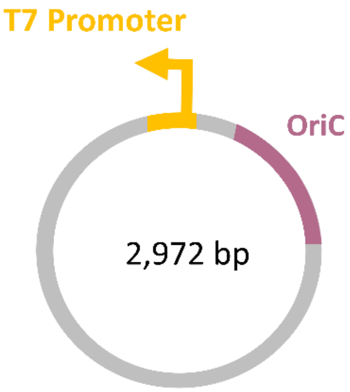


Table 5

| Sequence (5′→ 3′) |
|---|
| T7 promoter OriC ScaI cutting |
| TCGCGCGTTTCGGTGATGACGGTGAAAACCTCTGACACATGCAGCTCCCCTAGACGGTCACAGCTT |
| GTCTGTAAGCGGATGCCGGGAGCAGACAAGCCCGTCAGGGCGCGTCAGCGGGTGTTGGCGGGTGTC |
| GGGGCTGGCTTAACTATGCGGCATCAGAGCAGATTGTACTGAGAGTGCACCAAATGCGGTGTGAAA |
| TACCGCACAGATGCGTAAGGAGAAAATACCGCATCAGGCGCCATTCGCCATTCAGGCTGCGCAACT |
| GTTGGGAAGGGCGATCGGTGCGGGCCTCATCGCTATTACGCCAGCTGGCGAAAGGGGGATGTGCTG |
| CAAGGCGATTAAGTTGGGTAACGCCAGGGTTTTCCCAGTCACGACGTTGTAAAACGACGGCCAGTG |
| CAACGCGATGACGATGGATAGCGATTCATCGATGAGCTGACCCGATCGCCGCCGCCGGAGGGTTGC |
| GTTTGAGACAGGCGACAGATCCTGCAGGAAGGTTTAAACGCATTTAGGTGACACTATAGAAGTGGA |
| ATCCGCTCGAGGGATCCGAATTCGAAGCTTTGGTACAATTCGAGACCGGAGCGAGACGGGAGTCCA |
| GATCTCCATCGTCTCACCATGGTCTCAAGCTACCTGAAGCTTTCTTAATTAAGACGTCAGAATTCT |
| CGAGGCGGCCGCATGTGAGTCTC**CCTATAGTGAGTCGTATTA**ATCAGTTCTGGACCAGCGAGCTGT |
| GCTGCGACTCGTGGCGTAATCATGGTCATAGCTGTTTCCTGTGTGAAATTGTTATCCGCTCACAAT |
| TCCACACAACATACGAGCCGGAAGCATAAAGTGTAAAGCCTGGGGTGCCTAATGAGTGAGCTAACT |
| CACATTAATTGCGTTGCGCTCACTGCCCGCTTTCCAGTCGGGAAACCTGTCGTGCCAGCTGCATTA |
| ATGAATCGGCCAACGCGCGGGGAGAGGCGGTTTGCGTATTGGGCGCTCTTCCGCTTCCTCGCTCAC |
| TGACTCGCTGCGCTCGGTCGTTCGGCTGCGGCGAGCGGTATCAGCTCACTCAAAGGCGGTAATACG |
| GTTATCCACAGAATCAGGGGATAACGCAGGAAAGAACATGTGAGCAAAAGGCCAGCAAAAGGCCAG |
| GAACCGTAAAAAGGCCGCGTTGCTGGCGTT**TTTCCATAGGCTCCGCCCCCCTGACGAGCATCACAA** |
| **AAATCGACGCTCAAGTCAGAGGTGGCGAAACCCGACAGGACTATAAAGATACCAGGCGTTTCCCCC** |
| **TGGAAGCTCCCTCGTGCGCTCTCCTGTTCCGACCCTGTCGCTTACCGGATACCTGTCCGCCTTTCT** |
| **CCCTTCGGGAAGCGTGGCGCTTTCTCATAGCTCACGCTGTAGGTATCTCAGTTCGGTGTAGGTCGT** |
| **TCGCTCCAAGCTGGGCTGTGTGCACGAACCCCCCGTTCAGCCCGACCGCTGCGCCTTATCCGGTAA** |
| **CTATCGTCTTGAGTCCAACCCGGTAAGACACGACTTATCGCCACTGGCAGCAGCCACTGGTAACAG** |
| **GATTAGCAGAGCGAGGTATGTAGGCGGTGCTACAGAGTTCTTGAAGTGGTGGCCTAACTACGGCTA** |

**CACTAGAAGAACAGTATTTGGTATCTGCGCTCTGCTGAAGCCAGTTACCTTCGGAAAAAGAGTTGG**
**TAGCTCTTGATCCGGCAAACAAACCACCGCTGGTAGCGGTGGTTTTTTTGTTTGCAAGCAGCAGAT**
**TACGCGCAGAAAAAAAGGATCTCAA**GAAGATCCTTTGATCTTTTCTACGGGGTCTGACGCTCAGTG
GAACGAAAACTCACGTTAAGGGATTTTGGTCATGAGATTATCAAAAAGGATCTTCACCTAGATCCT
TTTAAATTAAAAATGAAGTTTTAAATCAATCTAAAGTATATATGAGTAAACTTGGTCTGACAGTTA
CCAATGCTTAATCAGTGAGGCACCTATCTCAGCGATCTGTCTATTTCGTTCATCCATAGTTGCCTG
ACTCCCCGTCGTGTAGATAACTACGATACGGGAGGGCTTACCATCTGGCCCCAGTGCTGCAATGAT
ACCGCGAGATCCACGCTCACCGGCTCCAGATTTATCAGCAATAAACCAGCCAGCCGGAAGGGCCGA
GCGCAGAAGTGGTCCTGCAACTTTATCCGCCTCCATCCAGTCTATTAATTGTTGCCGGGAAGCTAG
AGTAAGTAGTTCGCCAGTTAATAGTTTGCGCAACGTTGTTGCCATTGCTACAGGCATCGTGGTGTC
ACGCTCGTCGTTTGGTATGGCTTCATTCAGCTCCGGTTCCCAACGATCAAGGCGAGTTACATGATC
CCCCATGTTGTGCAAAAAAGCGGTTAGCTCCTTCGGTCCTCCGATCGTTGTCAGAAGTAAGTTGGC
CGCAGTGTTATCACTCATGGTTATGGCAGCACTGCATAATTCTCTTACTGTCATGCCATCCGTAAG
ATGCTTTTCTGTGACTGGTG**AGTACT**CAACCAAGTCATTCTGAGAATAGTGTATGCGGCGACCGAG
TTGCTCTTGCCCGGCGTCAATACGGGATAATACCGCGCCACATAGCAGAACTTTAAAAGTGCTCAT
CATTGGAAAACGTTCTTCGGGGCGAAAACTCTCAAGGATCTTACCGCTGTTGAGATCCAGTTCGAT
GTAACCCACTCGTGCACCCAACTGATCTTCAGCATCTTTTACTTTCACCAGCGTTTCTGGGTGAGC
AAAAACAGGAAGGCAAAATGCCGCAAAAAAGGGAATAAGGGCGACACGGAAATGTTGAATACTCAT
ACTCTACCTTTTTCAATATTATTGAAGCATTTATCAGGGTTATTGTCTCATGAGCGGATACATATT
TGAATGTATTTAGAAAAATAAACAAATAGGGGTTCCGCGCACATTTCCCCGAAAAGTGCCACCTGA
CGTCTAAGAAACCATTATTATCATGACATTAACCTATAAAAATAGGCGTATCACGAGGCCCTTTCA
TC

Map of the plasmid showing restriction sites, T7RNAP promoter sequence and the OriC.

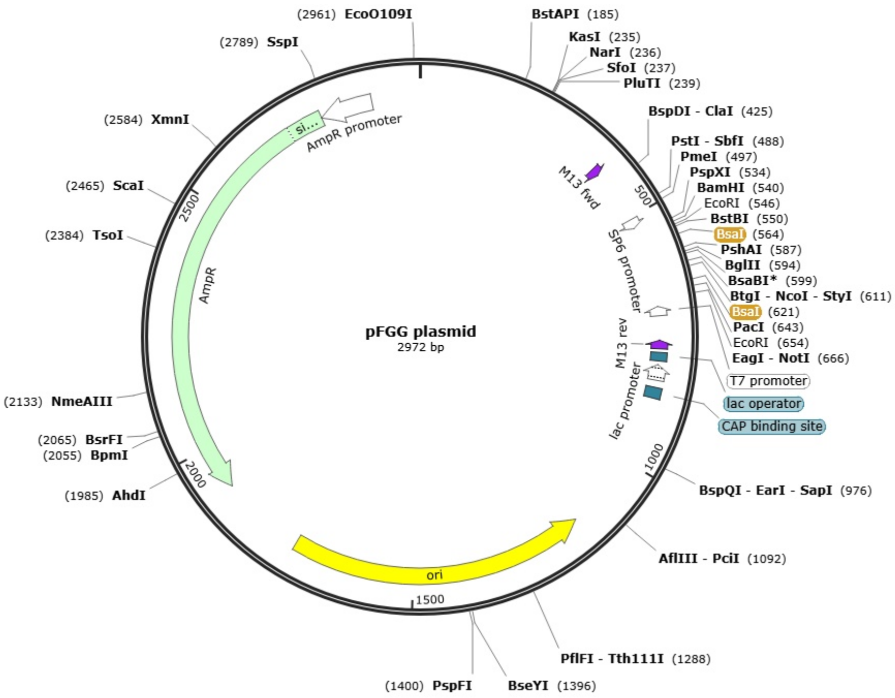

#### Table 6.

Sequence of oligonucleotides used to assemble RNA ID for characterization of transcripts originating from 4.5 kbp DNA construct. This construct has a larger separation between T7 promoter and OriC (Supplementary Table 4). This design is used for the experiments presented in Supplementary Figure 25.

Table 6

| Oligo number | Sequence (5′→ 3′) |
|---|---|
| 1 | AGCGTGATGCTACTAATTGGGACAATTTTCCAGATGAAGT |
| 2 | ATCATCTAAGAATTTAAATGAAGAAGACTTCAGAGCTTTT |
| 3 | GTTAAAAATTATTTGGCAAAAATAATATAATTCGGCTGCA |
| 4 | GGGGCGGCCTCGTGATACGCCTATTTTTATAGGTTAATGT |
| 5 | CATGATAATAATGGTTTCTTAGACGTCAGGTGGCACTTTT |
| 6 | CGGGGAAATGTGCGCGGAACCCCTATTTGTTTATTTTTCT |
| 7 | AAATACATTCAAATATGTATCCGCTCATGAGACAATAACC |
| 8 | CTGATAAATGCTTCAATAATATTGAAAAAGGAAGAGTATG |
| 9 | AGTATTCAACATTTCCGTGTCGCCCTTATTCCCTTTTTTG |
| 10 | CGGCATTTTGCCTTCCTGTTTTTAAGAAGGTTTGACCCAG |
| 11 | AAACGCTGGGAAAGGCGGTAAAATATGCACGAAGATCAGT |
| 12 | TCGGTTCACGAGTGGGTTACATCGAACTGGATCTCAACAG |
| 13 | CGGTAAGATCCTTGAAGAGTTTTCCGCCCCGAAGAACGTT |
| 14 | TTCCAATGATGAGCACTTTTAAAGCTCTGCTACTGTGGCG |
| 15 | CGGTTATATCCCGTATTGACGCCGGGCAAGAGCAACTCGG |
| 16 | TCGCCGCATACACTATTCTCAGAATGACTTGGTTGAGTAC |
| 17 | TCACCAGTCACAGAAAAGCATCTTACGGATGGCATGACAG |
| 18 | TAGAGAATTATGCAGTGCTGCCATAACCATGAGTGATAAC |
| 19 | ACTGCGGCCAACTTACTTCTGACAACGATCGGAGGACCGA |
| 20 | AGGAGCTAACCGCTTTTTTGCACAACATGGGGGATCATGT |
| 21 | AACTCGCCTTGATCGTTGGGAACCGGAGCTGAATGAAGCC |
| 22 | ATACCAAACGACGAGCGTGACACCACGATGCCTGTAGCAA |
| 23 | TGGCAACAACGTTGCGCAAACTATTAACTGGCGAACTACT |
| 24 | TACTCTAGCTTCCCGGCAACAATTAATAGACTGGATGGAG |
| 25 | GCGGATAAAGTTGCAGGACCACTTCTGCGCTCGGCCCTTC |
| 26 | CGGCTGGCTGGTTTATTGCTGATAAATCTGGAGCCGGTGA |
| 27 | GCGTGGGTCTCGCGGTATCATTGCAGCACTGGGGCCAGAT |
| 28 | GGTAAGCCCTCCCGTATCGTAGTTATCTACACGACGGGGA |
| 29 | GTCAGGCAACTATGGATGAACGAAATAGACAGATCGCTGA |
| 30 | GATAGGTGCCTCACTGATTAAGCATTGGTAACTGTCAGAC |
| 31 | CAAGTTTACTCATATATACTTTAGATTGATTTAAAACTTC |
| 32 | ATTTTTAATTTAAAAGGATCTAGGTGAAGATCCTTTTTGA |
| 33 | TAATCTCATGACCAAAATCCCTTAACGTGAGTTTTCGTTC |
| 34 | CACTGAGCGTCAGACCCCGTAGAAAAGATCAAAGGATCTT |
| 35 | CTTGAGATCCTTTTTTTCTGCGCGTAATCTGCTGCTTGCA |
| 36 | AACAAAAAAACCACCGCTACCAGCGGTGGTTTGTTTGCCG |

| | |
|---|---|
| 37 | GATCAAGAGCTACCAACTCTTTTTCCGAAGGTAACTGGCT |
| 38 | TCAGCAGAGCGCAGATACCAAATACTGTTCTTCTAGTGTA |
| 39 | GCCGTAGTTAGGCCACCACTTCAAGAACTCTGTAGCACCG |
| 40 | CCTACATACCTCGCTCTGCTAATCCTGTTACCAGTGGCTG |
| 41 | CTGCCAGTGGCGATAAGTCGTGTCTTACCGGGTTGGACTC |
| 42 | AAGACGATAGTTACCGGATAAGGCGCAGCGGTCGGGCTGA |
| 43 | ACGGGGGGTTCGTGCACACAGCCCAGCTTGGAGCGAACGA |
| 44 | CCTACACCGAACTGAGATACCTACAGCGTGAGCTATGAGA |
| 45 | AAGCGCCACGCTTCCCGAAGGGAGAAAGGCGGACAGGTAT |
| 46 | CCGGTAAGCGGCAGGGTCGGAACAGGAGAGCGCACGAGGG |
| 47 | AGCTTCCAGGGGGAAACGCCTGGTATCTTTATAGTCCTGT |
| 48 | CGGGTTTCGCCACCTCTGACTTGAGCGTCGATTTTTGTGA |
| 49 | TGCTCGTCAGGGGGGCGGAGCCTATGGAAAAACGCCAGCA |
| 50 | ACGCGGCCTTTTTACGGTTCCTGGCCTT |
| 51 | TTGCTGGCCTTTTGCTCACATGTTCTTTC |
| 52 | CTGCGTTATCCCCTGATTCTGTGGATTTGGATATCACTCATTAGTGGT |
| 53 | TAACCGTATTACCGCCTTTGAGTGAGCTGATACCGCTCGC |
| 54 | CGCAGCCGAACGACCGAGCGCAGCGAGTCAGTGAGCGAGG |
| 55 | AAGCGGAAGAGCGCCCAATACGCAAACCGCCTCTCCCCGC |
| 56 | GCGTTGGCCGATTCATTAATGCAGCTGGCACGACAGGTTT |
| 57 | CCCGACTGGAAAGCAATTGGCAGTGAGCGCAACGCAATTA |
| 58 | ATGTGAGTTAGCTCACTCATTAGGCACCCCAGGCTTTACA |
| 59 | CTTTATGCTTCCGGCTCGTATAATGTGCTGATGAATCCCC |
| 60 | TAATGATTTTGGTAAAAATCATTAAGTTAAGGTGGATACA |
| 61 | CATCTTGTCATATGATCAAATGGTTTCGCGAAAAATCAAT |
| 62 | AATCAGACAACAAGATGTGCGAACTCGATATTTTACACGA |
| 63 | CTCTCTTTACCAATTCTGCCCCGAATTACACTTAAAACGA |
| 64 | CTCAACAGCTTAACGTTGGCTTGCCACGCATTACTTGACT |
| 65 | GTAAAACTCTCACTCTTACCGAACTTGGCCGTAACCTGCC |
| 66 | AACCAAAGCGAGAACAAAACATAACATCAAACGAATCGAC |
| 67 | CGATTGTTAGGTAATCGTCACCTCCACAAAGAGCGACTCG |
| 68 | CTGTATACCGTTGGCATGCTAGCTTTATCTGTTCGGGCAA |
| 69 | TACGATGCCCATTGTACTTGTTGACTGGTCTGATATTCGT |
| 70 | GAGCAAAAACGACTTATGGTATTGCGAGCTTCAGTCGCAC |
| 71 | TACACGGTCGTTCTGTTACTCTTTATGAGAAAGCGTTCCC |
| 72 | GCTTTCAGAGCAATGTTCAAAGAAAGCTCATGACCAATTT |
| 73 | CTAGCCGACCTTGCGAGCATTCTACCGAGTAACACCACAC |
| 74 | CGCTCATTGTCAGTGATGCTGGCTTTAAAGTGCCA |
| 75 | TGGTATAAATCCGTTGAGAAGCTGGTTTGGATATCACTCATTAGTGGT |
| 76 | GTTGGTACTGGTTAAGTCGAGTAAGAGGAAAAGTACAATA |
| 77 | TGCAGACCTAGGAGCGGAAAAACTGGAAACCTATCAGCAA |
| 78 | CTTACATGATATGTCATCTAGTCACTCAAAGACTTTAGGC |
| 79 | TATAAGAGGCTGACTAAAAGCAATCCAATCTCATGCCAAA |
| 80 | TTCTATTGTATAAATCTCGCTCTAAAGGCCGAAAAAATCA |
| 81 | GCGCTCGACACGGACTCATTGTCACCACCCGTCACCTAAA |
| 82 | ATCTACTCAGCGTCGGCAAAGGAGCCATGGGTTCTAGCAA |

| | |
|---|---|
| 83 | CTAACTTACCTGTTGAAATTCGAACACCCAAACAACTTGT |
| 84 | TAATATCTATTCGAAGCGAATGCAGATTGAAGAAACCTTC |
| 85 | CGAGACTTGAAAAGTCCTGCCTACGGACTAGGCCTACGCC |
| 86 | ATAGCCGAACGAGCAGCTCAGAGCGTTTTGATATCATGCT |
| 87 | GCTAATCGCCCTGATGCTTCAACTAACATGTTGGCTTGCG |
| 88 | GGCGTTCATGCTCAGAAACAAGGTTGGGACAAGCACTTCC |
| 89 | AGGCTAACACAGTCAGAAATCGAAACGTACTCTCAACAGT |
| 90 | TCGCTTAGGCATGGAAGTTTTGCGGCATTCTGGCTACACA |
| 91 | ATAACAAGGGAAGACTTACTCGTGGCTGCAACCCTACTAG |
| 92 | CTCAAAATTTATTCACACATGGTTACGCTTTGGGGAAATT |
| 93 | ATGAGGGGATCTCTCAGTATAATGTGTGGAATTATGAGCG |
| 94 | GATAATAATTTCACACAGGAGGTTTAAACTTTAAACATGT |
| 95 | CAAAAGAGACGTCTTTTGTTAAGAATGCTGAGGAACTTGC |
| 96 | AAAGCAAAAAAATGGATGCTATTAACCCTGAACTTTCTTCA |
| 97 | AAATTTAAATTTTTAATAAAATTCCTGTCTCAGTT |
| 98 | TCCTGAAGCTTGCTCTAAACCTCGTTTTGGATATCACTCATTAGTGGT |
| 99 | TCAAAAAAAATGCAGAATAAAGTTGTTTGGATATCACTCATTAGTGGT |
| 100 | GTCAAGAGGAACATATTGAATATTTAGCTCGTAGTTTTCA |
| 101 | TGAGAGTCGATTGCCAAGAAAACCCACGCCACCTACAACG |
| 102 | GTTCCTGATGAGGTGGTTAGCATAGTTCTTAATATAAGTT |
| 103 | TTAATATACAGCCTGAAAATCTTGAGAGAATAAAAGAAGA |
| 104 | ACATCGATTTTCCATGGCAGCTGAGAATATTGTAGGAGAT |
| 105 | CTTCTAGAAAGATTTAAGCCG |
| 106 | AGAATGGTCTGTGATCCCCCC |
| 107 | CATTCCCGGCTACACTGCACCATGATCTTGCTGAAAAACT |
| 108 | CGAGCCATCCGGAAGATCTGGCGGCCGCTCTCCC |
| 109 | CAGCAGCAGCAGCAGCAGCAGCAGCAGCAG |

#### Table 7.

Identified sequence within DNA construct that shows structural similarity to engineered T7RNAP transcription terminators.

Table 7

| DNA | Sequence (5′→ 3′) |
|---|---|
| 1 | CAAACAAACCACCGCTGGTAGCGGTGGTTTTTTTGTTT |

**Table 8.**

The table shows the IV curves of nanopores used to study RNA IDs. The ionic current and RMS noise at 600 mV is presented for each pore. An estimate of each nanopore diameter was calculated from the ionic current values presented, assuming a conical pore geometry[5].

Table 8

| **Pore number** | **Sample** | **Current, RMS noise at 600 mV and calculated pore diameter** | **Current / voltage curve (IV curve)** |
|---|---|---|---|
| 1 | RNA IDs produced from linear DNA | 11.4 nA<br>6.5 pA<br>~8 nm | 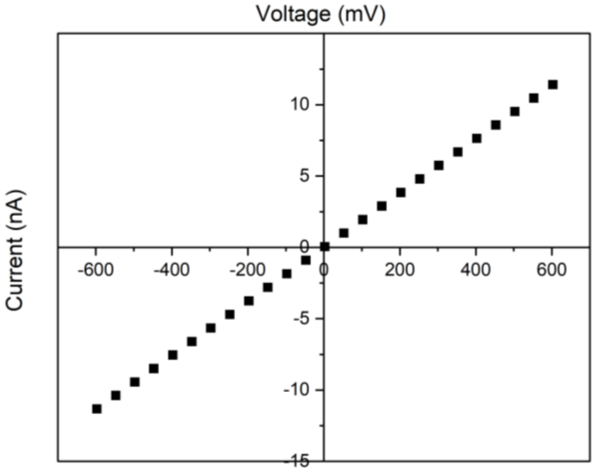 |
| 2 | RNA IDs produced from linear DNA, and DNA template | 9.9 nA<br>6.7 pA<br>~7 nm | 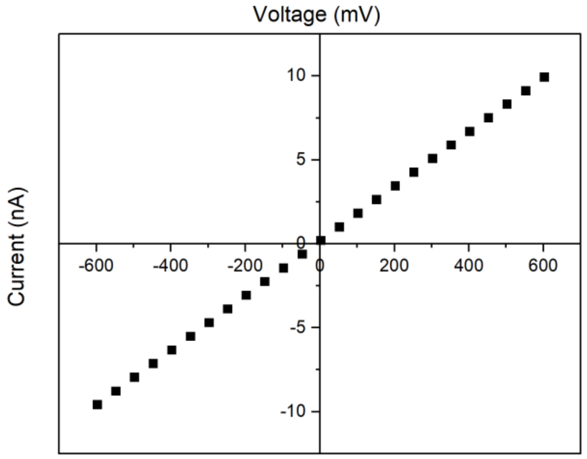 |

| | | | |
|---|---|---|---|
| 3 | RNA IDs of produced from circular template | 8.7 nA<br>6.2 pA<br>~6 nm | 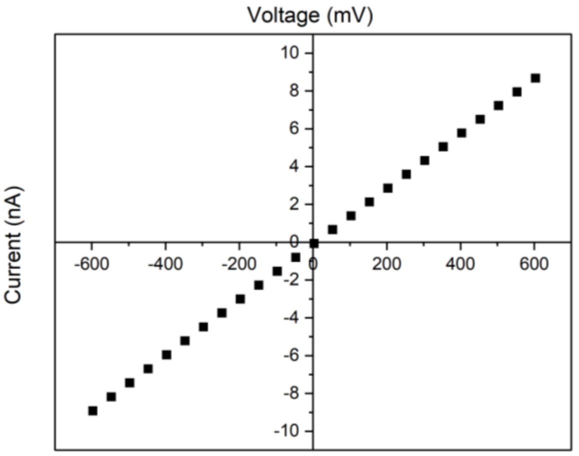 |
| 4 | RNA IDs of produced from circular template | 7.5 nA<br>6.6 pA<br>~5 nm | 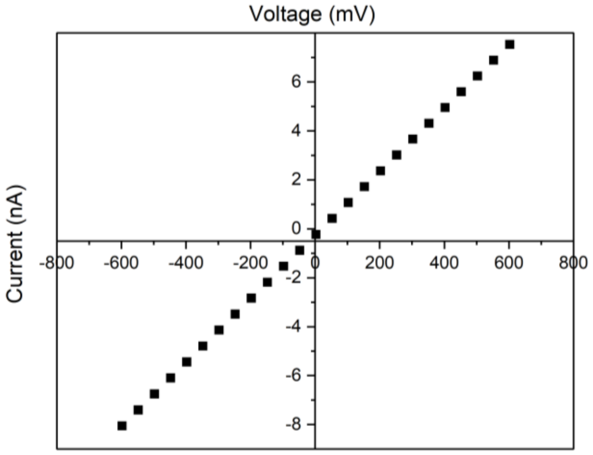 |
| 5 | RNA IDs of produced from circular template | 12.2 nA<br>6.4 pA<br>~9 nm | 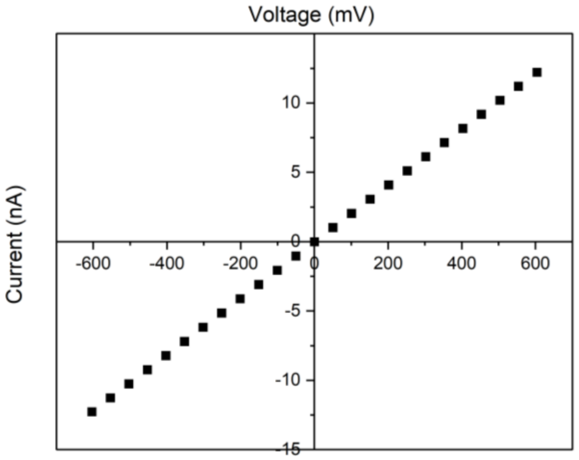 |

| | | | |
|---|---|---|---|
| 6 | RNA IDs of produced from circular template | 10.0 nA<br>6.4 pA<br>~7 nm | 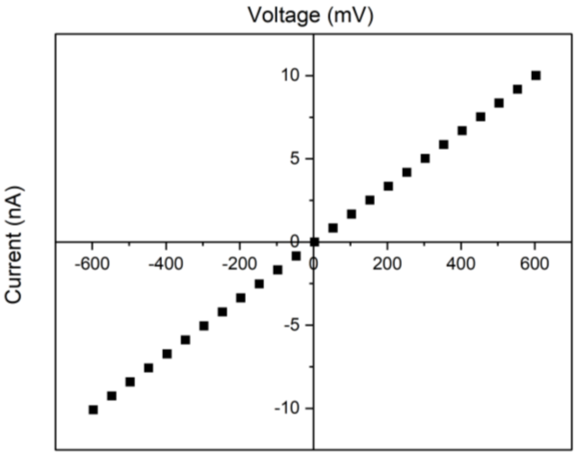 |
| 7 | DNA ladder SF8 (0.5 kbp – 10 kbp) | 11.1 nA<br>5.7 pA<br>~8 nm | 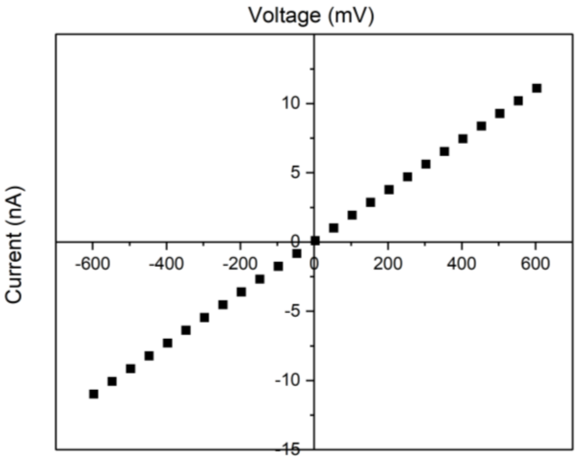 |
| 8 | RNA IDs produced from 4.5 kbp DNA | 10.9 nA<br>6.4 pA<br>~8 nm | 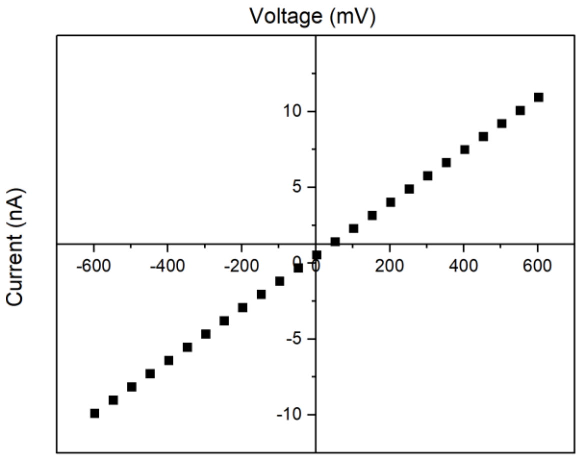 |

An estimate the pore's diameter was calculated from the overall resistance of the nanopore in open state, R, which is constituted by the resistance of the pore cavity region, $R_{pore}$, and the resistance of the access region of the pore, $R_{acc}$[5]:

$$R = R_{pore} + R_{acc}$$

This equation can be rewritten in terms of the resistivity $\rho$ of the electrolytic solution, the pore's length, $L$, and the diameter of the *cis* and *trans* aperture of the pore, $D_{cis}$ and $D_{trans}$ :

$$R = \rho \frac{4L}{\pi D_{trans} D_{cis}} + \rho \left(\frac{1}{2 D_{trans}} + \frac{1}{2 D_{cis}}\right)$$

The diameter of the pore $D_{cis}$ was calculated using the experimental ionic current $I$ during the application of a 600 mV potential, assuming a $D_{trans}$ of 200 µm, conductivity of $15.5\ Sm^{-1}$ for 4M LiCl, and length $L$ of 950 µm for our glass nanopores [6,7].
